# Mismatch responses mediated by adaptation and deviance detection have complementary functional profiles that point to different auditory short-term memory systems

**DOI:** 10.1101/2019.12.18.881821

**Authors:** Tobias Teichert, Hank Jedema, Zhijun Shen, Kate Gurnsey

**Affiliations:** Department of Psychiatry, University of Pittsburgh, Pittsburgh, PA; Department of Bioengineering, University of Pittsburgh, Pittsburgh, PA; Preclinical Pharmacology Section, NIDA, Baltimore, MD

## Abstract

Mismatch negativity (MMN) is a macroscopic EEG deflection in response to rare or unexpected sounds. It has provided important insights into auditory short-term memory, pre-attentive guidance of attention, and their alteration in conditions such as schizophrenia. It remains unclear if MMN is caused by passive adaptation, active memory-comparison processes (deviance detection; DD), or a mix of both. To answer this question, macaque monkeys listened to a new paradigm that quantified both components of MMN. Micro- and macroscopic mismatch responses in the rhesus were dominated by adaptation at short latencies but included a smaller contribution of deviance detection at longer latencies. Most importantly, we show that mismatch responses mediated by adaptation have a short temporal scope and narrow frequency tuning while mismatch responses mediated by deviance detection have a longer temporal scope but broader frequency tuning. The different functional profiles point to the involvement of two distinct auditory short-term memory systems and complementary roles in the pre-attentive guidance of attention.

## 1 Background

Mismatch negativity (**MMN**) is a fronto-central, surface-negative EEG deflection in response to a rare or unexpected auditory event (Näätänen and Picton, 1987). Due to its non-invasive nature, MMN can readily be recorded in the human, where it has provided important insights into auditory short-term memory (Näätänen et al., 2005) and pre-attentive neural mechanisms that help prioritize processing of potentially informative events (Jääskeläinen et al., 2004). In addition to elucidating normal brain function, MMN is also a sensitive and selective marker of altered auditory function and progressive auditory cortex gray matter loss in schizophrenia (**SZ**)(Javitt and Sweet, 2015; Salisbury et al., 2007). Despite its importance, it is still unclear what the macroscopic MMN reveals about the underlying neural computations. Determining this relationship would pave the way for a more detailed inference of disease-relevant neural mechanisms from non-invasive measurements at the scalp. In particular, there is an ongoing debate about whether the MMN largely reflects (a) release from adaptation (Jääskeläinen et al., 2004; May and Tiitinen, 2010; Ulanovsky et al., 2003), i.e., a relatively simple passive process that can be implemented at the level of individual synapses and/or neurons or (b) an active memory-comparison processes (Näätänen et al., 2005), i.e., a rather complex computation as recently elaborated in the predictive coding framework (Friston, 2005; Lieder et al., 2013; Rao and Ballard, 1999; Wacongne et al., 2012), which envisions sophisticated cascades of feedforward and feedback projections between multiple processing stages.

Adaptation and predictive coding themselves can be broken down into distinct components, and a detailed understanding of the MMN necessarily requires a detailed understanding of these components and their interactions. Adaptation can be divided into three distinct phenomenological components that describe reductions of neural responses in three different scenarios (Figure 1A). The most well-known component is *stimulus-specific* adaptation (Antunes and Malmierca, 2014; Nieto-Diego and Malmierca, 2016; Ulanovsky et al., 2004, 2003)(**SSA**) which describes reductions of neural responses as a function of the proximity of preceding stimuli in feature space: adaptation is stronger when the stimuli are identical or very similar. Adaptation also has a prominent temporal component (refractoriness, **R**) which describes reductions of neural responses as a function of the proximity of preceding stimuli in time (Budd et al., 1998; Lu et al., 1992a, 1992b; Pereira et al., 2014; Ritter et al., 1968): adaptation is stronger when the auditory system is given less time to recover between sounds. Finally, *short-term habituation* (**STH**) or *repetition suppression* describes reductions of neural responses as a function of how often the same tone has been repeated (Budd et al., 1998; Callaway, 1973; Loveless, 1983): adaptation is stronger when a sound has been repeated more often. STH provides insight into how the brain integrates the effects of more than one adapting event. In the context of MMN, weaker responses to the standards and stronger responses to the deviants may arise because the deviants are presented at a slower rate (less R), are not repeated as often as the standards (less STH) and are physically distinct from the preceding standards (less SSA).

**Figure 1.**
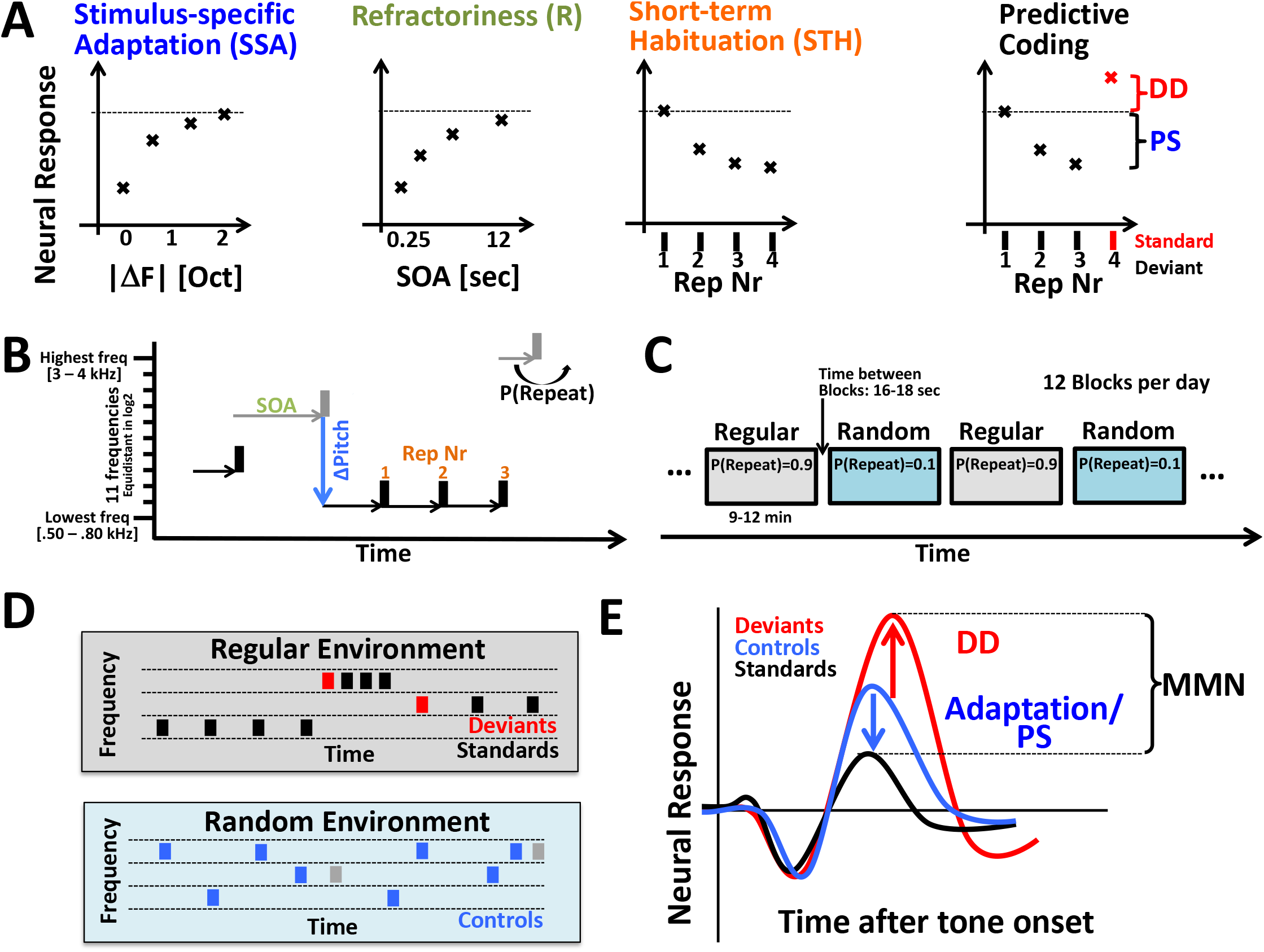
Methods. **(A)** Theoretical breakdown of MMN into adaptation (SSA, R, STH) and predictive processing (PS and DD). SSA: neural response to the second of two tones decreases if the two tones have a smaller difference in fundamental frequency. R: neural responses to the second of two tones decreases if the preceding tone was closer in time. STH: neural responses decrease with repetition of the same tone. The predictive coding account of MMN postulates two components: Predictive suppression (PS) attenuates responses to expected tones. Deviance detection (DD) enhances neural response to a tone it violates an explicit prediction. MMN may reflect a mix of all four of these components. (**B**) The RS-kernel paradigm is a modified roving standard paradigm that varies not only pitch, but also SOA and the probability that a combination of SOA and pitch is repeated (P(Repeat)). If P(Repeat) is high (low), the auditory environment is (ir)regular and (un)predictable. (**C**) Sound sequences are presented in either regular or irregular blocks of 8-12 minutes duration. (**D**) Example sequences of tones in the regular and random environment. The regular environment gives rise to deviants (red) and standards (black). The random environment gives rise to controls (blue). (**E**) MMN is defined as the difference between all deviants and all controls, regardless of SOA and repetition number. MMN is broken down into adaptation/PS, the difference between the controls and the standards, and DD, the difference between the deviants and the controls.

Predictive coding holds that the auditory system extracts regularities from the environment and uses these regularities to make explicit predictions about upcoming sounds. When an unpredicted stimulus, i.e., a deviant, is presented, it triggers stronger neural responses which may reflect the explicit detection of an unexpected deviant (*deviance detection*, DD), the mismatch between the expected sound and the deviant (*prediction error*), or additional computations to update the brain’s model of the auditory environment (*model update*). While these interpretations differ conceptually (Lieder et al., 2013), they make similar predictions in the current context. We will thus use the term DD as a placeholder for all of these active processes. DD has been suggested to account for MMN in situations that are difficult to explain with adaptation, such as unexpected stimulus omissions, or unexpected stimulus repetitions.

Several groups have developed novel experimental approaches such as the ‘*many-standards control condition*’ to identify and quantify contributions of adaptation and DD to the MMN (Schröger and Wolff, 1996). The many-standards control condition is comprised of control tones which presumably match the level of adaptation of the deviants, but do not violate an established auditory regularity. Stronger responses to the deviants compared to the controls would thus reflect an active process such as DD rather than adaptation (Figure 1E). Conversely, weaker responses to the standards compared to the controls would be consistent with adaptation. Indeed, the control condition has been found to split the MMN into two components (Jacobsen and Schröger, 2001) that emerge at slightly different latencies and have been localized to slightly different generators within auditory cortex (Maess et al., 2007; Opitz et al., 2005). However, other than these relatively subtle differences, the two putative components have remained somewhat amorphous and a clear functional distinction between them has yet to come into focus. This has given rise to the concern that the dissection of the MMN into SSA and DD may not reflect the existence of two distinct mechanisms, but rather a failure of the control condition to appropriately match the level of adaptation of the deviants. In summary, it remains entirely possible that the MMN is mediated either by release from adaptation, predictive coding or some mixture of the two.

The uncertainty about whether the segregation of the MMN into adaptation and DD is artificial or reflective of two distinct neural mechanisms may have slowed efforts to search for meaningful functional differences between the two. It seems likely that the evolution of two distinct mechanisms would be facilitated if both provide meaningfully distinct functionality. For example, the auditory system recovers from adaptation rather quickly, thus limiting the temporal scope of any function mediated by adaptation. To date, however, it is not known if DD could provide added functionality in the form of a longer life-time. Similarly, it is possible that the versatility of a mechanism such as DD comes at some cost such as reduced stimulus sensitivity. To date, however, it is unknown if SSA and DD are equally sensitive to physical differences between standards and deviants. The presence (or absence) of such functional distinctions between SSA and DD would then inform the original question about one or two neural mechanisms contributing to the MMN.

To approach this issue empirically, we modified the commonly used roving-standard paradigm (Baldeweg et al., 2004; Christianson et al., 2014; Costa-Faidella et al., 2011; Garrido et al., 2008; Lieder et al., 2013; Schröger and Wolff, 1996; Teichert et al., 2016) in a way that allowed us to measure key functional properties of the presumed mechanisms contributing to the MMN such as their latency, life-time, and frequency tuning. The experiments were conducted in non-human primates whose MMN is similar to the human with respect to timing, polarity, and a topography that points to a putative neural generator in auditory cortex (Gil-da-Costa et al., 2013; Honing et al., 2012). Using the monkey allowed us to record not only the macroscopic MMN using EEG, but also established microscopic mismatch responses (MMR) in auditory cortex (Camalier et al., 2019; Fishman, 2014; Fishman and Steinschneider, 2012; Javitt et al., 1996, 1992; Lakatos et al., 2019), and to determine whether different populations of cells preferentially encode aspects of adaptation, predictive coding or both. Building on earlier work (Christianson et al., 2014; Herrmann et al., 2016; Taaseh et al., 2011; Teichert et al., 2016), we developed a detailed feed-forward computational model of adaptation based on short-term synaptic depression at thalamo-cortical synapses. The computational model provides an objective test of the core assumption that adaptation in the many-standard control condition matched the level of adaptation of the deviants.

Briefly, our results confirm that we can reliably decompose the mismatch response into two components with clearly distinct functional properties. The presumed adaptation component accounts for the bulk of the mismatch response in the early stimulus-evoked period and has relatively sharp frequency tuning but a relatively short life-time. In contrast, the putative contribution of DD emerges later, requires larger frequency differences but is more stable if tones are spaced out farther in time. Taken together, our findings suggest that the first wave of mismatch responses in auditory cortex is dominated by stimulus-specific and non-specific adaptation, while a second wave is at least in part driven by DD, which may be fed back from higher areas. The data suggests that passive adaptation and active prediction both contribute to the MMN and may play distinct yet complementary roles in prioritizing informative sounds.

## 2 Results

### 2.1 Stimulus-specific adaptation and deviance detection in macroscopic EEG responses

To characterize mismatch responses at the macroscopic level, we recorded EEG activity in four male macaque monkeys from up to 33 chronically implanted cranial EEG electrodes that were embedded in 1mm deep holes in the skull and covered with dental acrylic (Teichert, 2016). Electrodes were positioned in a grid-like arrangement that provided approximately equidistant electrode spacing across the accessible part of the skull, similar to the international 10-20 system in the human (Li and Teichert, 2020). EEG activity was recorded while animals passively listened to a modified roving standard paradigm (Figure 1B-D). In the regular condition, the subjects passively listened to pure tones that were predictable in time and tonal frequency (‘*standard*’). For 10% of the tones, the predictable sequence was disrupted by a tone that established a new temporal and tonal pattern (‘*deviant*’). In 8-12 minute long alternating blocks, tones were presented without regularity or deviations thereof and these tones served as a control for the lower degree of (stimulus specific) adaptation of deviants compared to standards (‘*control*’). Following conventions, MMN, or, more generally speaking, MMR was defined as the difference between the deviants and the standards. The MMR was broken up into SSA, which was operationalized as the difference between the controls and the standards, and DD which was operationalized as the difference between deviants and controls.

Figure 2A shows the average evoked responses at electrode Fz to the standard (red) and the deviant (black) across all sessions in example animal S. The two traces begin to diverge around the time of the P21 giving rise to a mismatch positivity that extends until around 65 ms after stimulus onset. Subsequently, the sign of the difference inverts giving rise to the classical MMN that peaks after the peak of the N85, the presumed N1 homolog of the monkey. Figure 2B plots the difference between the deviants and standards thus isolating the timing and significance of the MMR, again highlighting the biphasic pattern that features an early mismatch positivity followed by a later mismatch negativity. Both of these components reach a statistical criterion of α<0.001 in a time-resolved t-test. A second animal very closely matched this response pattern (animal J). A third animal exhibited only the mismatch positivity (animal W), and the fourth animal only exhibited the mismatch negativity (animal R). Despite these clear inter-individual differences, the data provide solid evidence for a previously unidentified mismatch positivity in 3 out of 4 animals, and confirms the previously reported mismatch negativity, also in 3 out of 4 animals.

**Figure 2.**
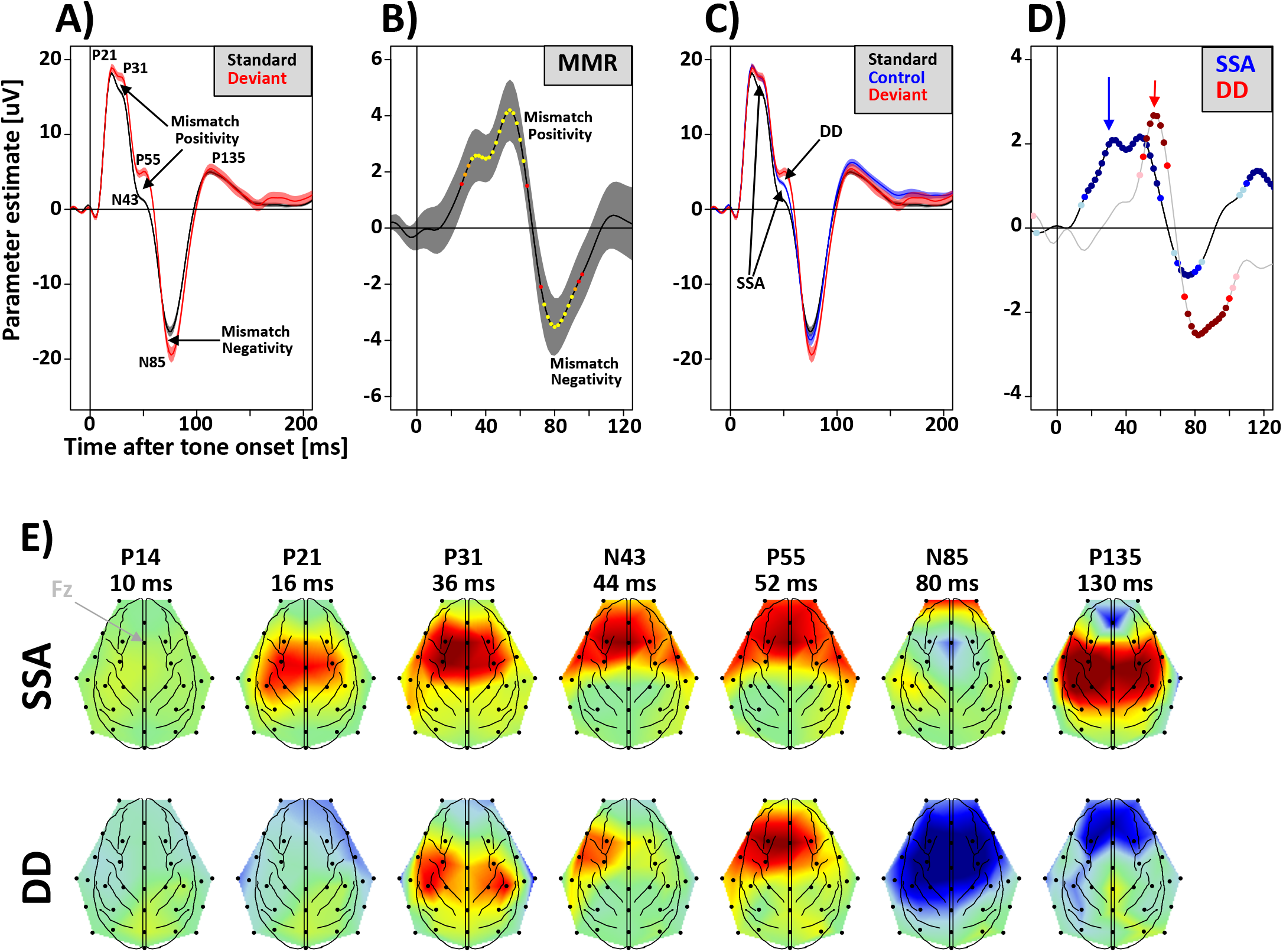
EEG mismatch responses in one example animal. (A) Responses to the standard (black) and the deviant (red) as a function of time from tone onset at electrode Fz. Error bars represent the standard error of the mean. Peaks and troughs are labeled based on our earlier classification scheme (Teichert et al. 2016a). (B) Mismatch responses quantified as the difference between the deviant and the standard. Dots of different color indicate p-value of a running t-test (red: p<.05; orange: p<.01; red: p<.001). (C) Data from the many-standards control condition is used to segregate mismatch responses into stimulus-specific adaptation (SSA) and deviance detection (DD). (D) SSA is quantified as the difference between the control and the standard (blue); DD is defined as the difference between the deviant and the control (red). Dots of different saturation indicate p-value of a running t-test (light blue/red: p<.05; blue/red: p<.01; dark blue/red: p<.001 (E) Topography of SSA and DD at the average latency of 7 previously identified EEG components (P14, P21, …). SSA emerges around the time of the P21 which has an average latency of 16 ms in this animal. During the time of the P31, DD starts emerging over fronto-lateral electrodes. DD peaks around the time of the P55.

Figure 2C uses the controls (blue) to segregate the MMRs into components related to SSA and DD, respectively. If the entire MMR were to be accounted for by SSA, the controls would overlap with the deviants. If the entire MMR were to be accounted for by DD, the controls would overlap with the standards. Data in this example show that early on, around the time of the P21 and P31, the MMR is explained exclusively by SSA. Later on, however, around the time of the P55, the control condition falls between the standard and the deviant, thus suggesting that the MMR is driven by both SSA and DD. Figure 2D isolates the SSA and DD component of the MMR to provide a clearer look at the timing and significance of the two components. The data confirm that significant SSA emerges first, at around 20 ms after tone-onset. DD emerges later, peaking around 55 ms after tone onset. During the time of the classical MMN, both SSA and DD are present. Figure 2E displays the topography of SSA and DD around the time of 7 previously identified EEG components (P14, P21, P31, N43, P55, N85 and P135). SSA emerges around the time of the P21 with a fronto-central topography that seems to shift more anterior around the time of the P31, N43 and P55. DD first emerges around the time of the P31 over centro-lateral electrodes. Around the time of the P55 and N85, DD exhibits the classic fronto-central topography.

Data from the other animals reveals clear inter-individual differences. However, there are several important commonalities across animals: (a) In 3 out of 4 animals the P21 component is significantly reduced by SSA; (b) In 2 of 3 animals that exhibit a P31, it is significantly reduced by SSA; (c) In 3 out of 4 animals the P55 is significantly enhanced by DD; (d) In all animals that show evidence of both components, SSA emerges earlier than DD.

The composition of the MMR around the time of the N85, i.e., the classical MMN, is less consistent across animals. Two of the 3 animals that have an MMN, also show evidence of the SSA subcomponent (animals S and R). Only one of them (animal S) shows evidence of a DD subcomponent. In the third animal with an MMN (animal J) neither SSA nor DD reach significance. However, it is worth noting two potential reasons. 1) The MMN in this animal is split between SSA and DD almost equally, thus reducing the likelihood of either component reaching significance. 2) Both subcomponents are prolonged and rather large (~2uV), suggesting that it the lack of a significant effect may be driven by large variability, rather than small difference in the means.

Figure 3 depicts the grand average over all four subjects using the same format as Figure 2A-D. The grand averages recapitulate the key findings identified in example animal in Figure 2 including an early mismatch positivity followed by a mismatch negativity that peaks shortly after the peak of the N85. Furthermore, there is evidence for the existence of SSA as well as DD, and that the SSA emerges earlier than DD. While key effects reach significance, it is important not to over-interpret these population statistics. First, there is a good bit of inter-subject variability regarding existence and timing of different components (see above). For example, animal W does not exhibit the P31, a key feature of the other three animals. Second, the power of these tests with an N of 4 is rather low. Typically, they reach significance only if all four subjects show the same effect at comparable magnitudes. Significance is lost not only if the effect is particularly small or absent in one animal, but also if the effect is particularly large in one animal, thus increasing its variance.

**Figure 3.**
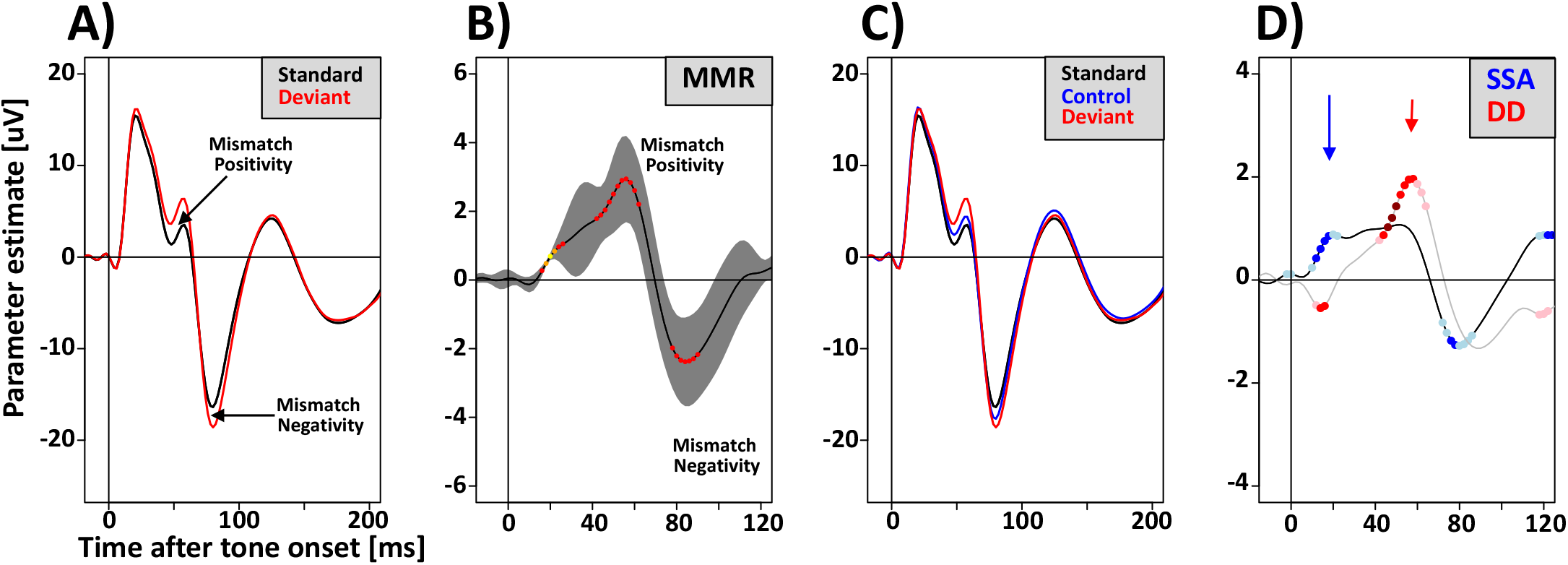
Population EEG mismatch responses across all four animals. Conventions as in Figure 2A-D. Error bars in B represent standard error of the means across the 4 animals.

### 2.2 Stimulus-specific adaptation and deviance detection in A1 multi-unit activity

To characterize mismatch responses at the microscopic level, we implanted a recording chamber over the left hemisphere to provide approximately perpendicular access to auditory cortex in the superior temporal plane in two of the animals (animals S and W). Early on, recordings were made through small burr-holes, leaving all EEG electrodes intact. Later on, the skull inside the chamber, and with it one of the EEG electrodes were removed.

Figure 4 shows the average laminar response profile for multi-unit activity, local field potentials and current source densities averaged across all sessions in animal S. The putative cortical layer of each recording site on the probe was estimated based on the location of the prominent supra-granular source-sink pair described previously. We identified a multi-unit as tone-responsive if the time-resolved linear model contained at least 5 time-bins in which the intercept was significant at a Bonferroni-Holmes corrected alpha-value of 0.01.

**Figure 4.**
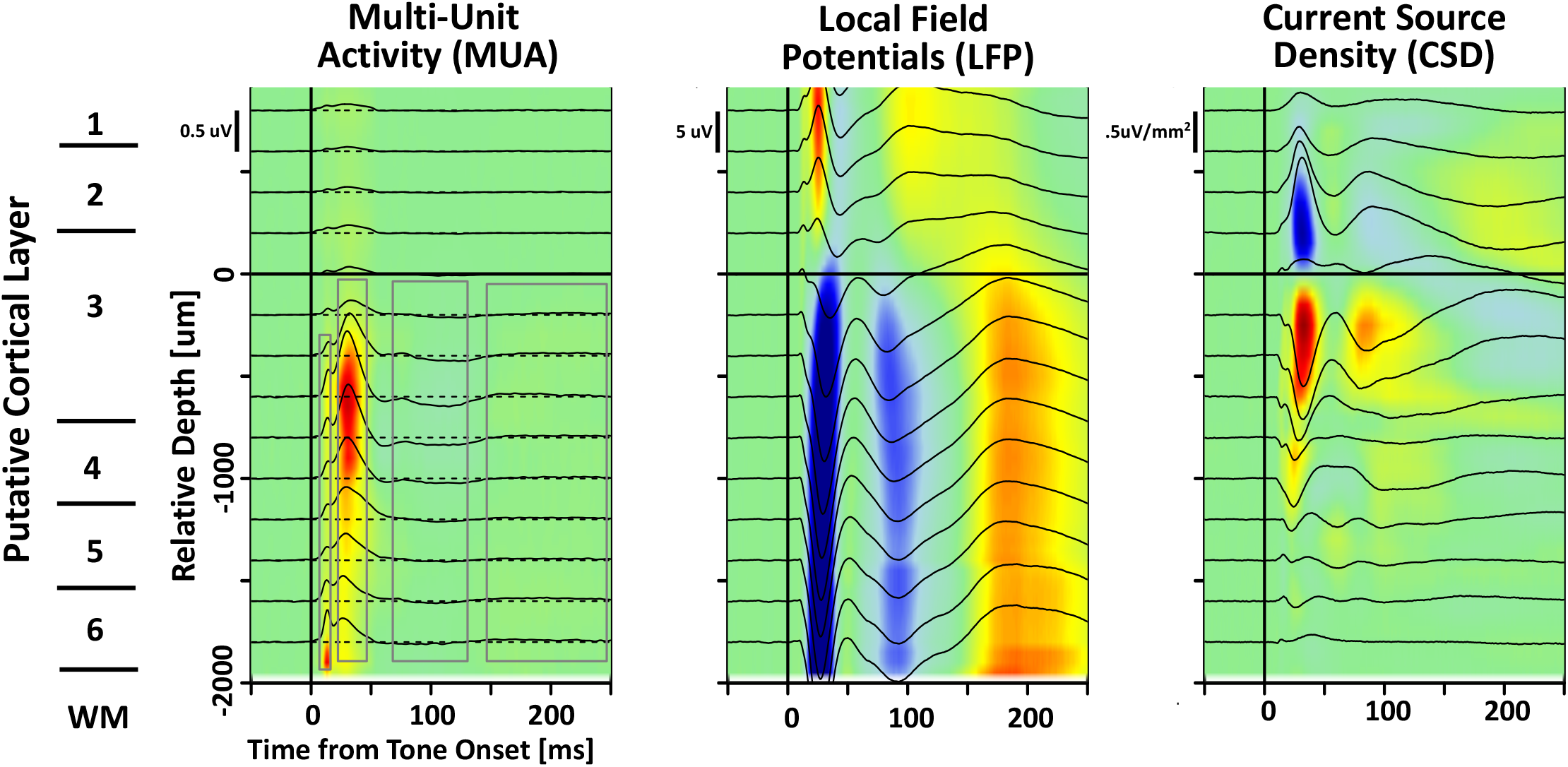
Average MUA, LFP and CSD profile in one example animal. Neural responses as a function of time from tone onset (x-axis) and putative cortical depth (y-axis). Different panels highlight different aspects of neural activity: multi-unit activity (left panel), local field potentials (middle panel), and current source density (right panel). Putative cortical depth was assigned based on the location of the supra-granular source-sink pair described in the literature. Multi-unit activity was divided into four main time-bins: (i) an early wave of activity likely corresponding to afferent thalamic fibers with a latency of 8 – 12 ms (th); the first wave of cortical evoked activity with a latency of 25 to 45 ms (evk); a period of suppressed activity between 70 and 120 ms (sup); and a period of rebound activity from 150 to 250 ms (reb). Based on the known anatomical extent of thalamic innervation that was confirmed by the functional measurements here, the thalamic epoch only included contacts with a relative depth below deep layer 3 (< −500 um).

Figure 5A&C plots the average activity of all tone-responsive multi-units in response to the standards, controls and deviants in both animals. The early peak around 12 ms after tone-onset presumably corresponds to the activity in afferent thalamic fibers. At the time of the peak of the multi-unit response (~30 ms), the standard elicits weaker responses than the control and deviant. After around 50 ms, all three conditions begin to separate. The strongest responses are observed for the deviant, followed by the control and the standard. Figure 5B&D isolates the SSA and DD component of the MMR to provide a closer look at the timing and significance of the two components. Both animals show clear SSA that starts emerging immediately after the peak of the presumed thalamic fiber activity at around 12 ms after tone onset. SSA reaches its maximum around 35 ms, briefly after the peak in cortical multi-unit activity. Significant DD is also present in both animals but reaches its peak on the falling slope of the cortical multi-unit activity and SSA.

**Figure 5.**
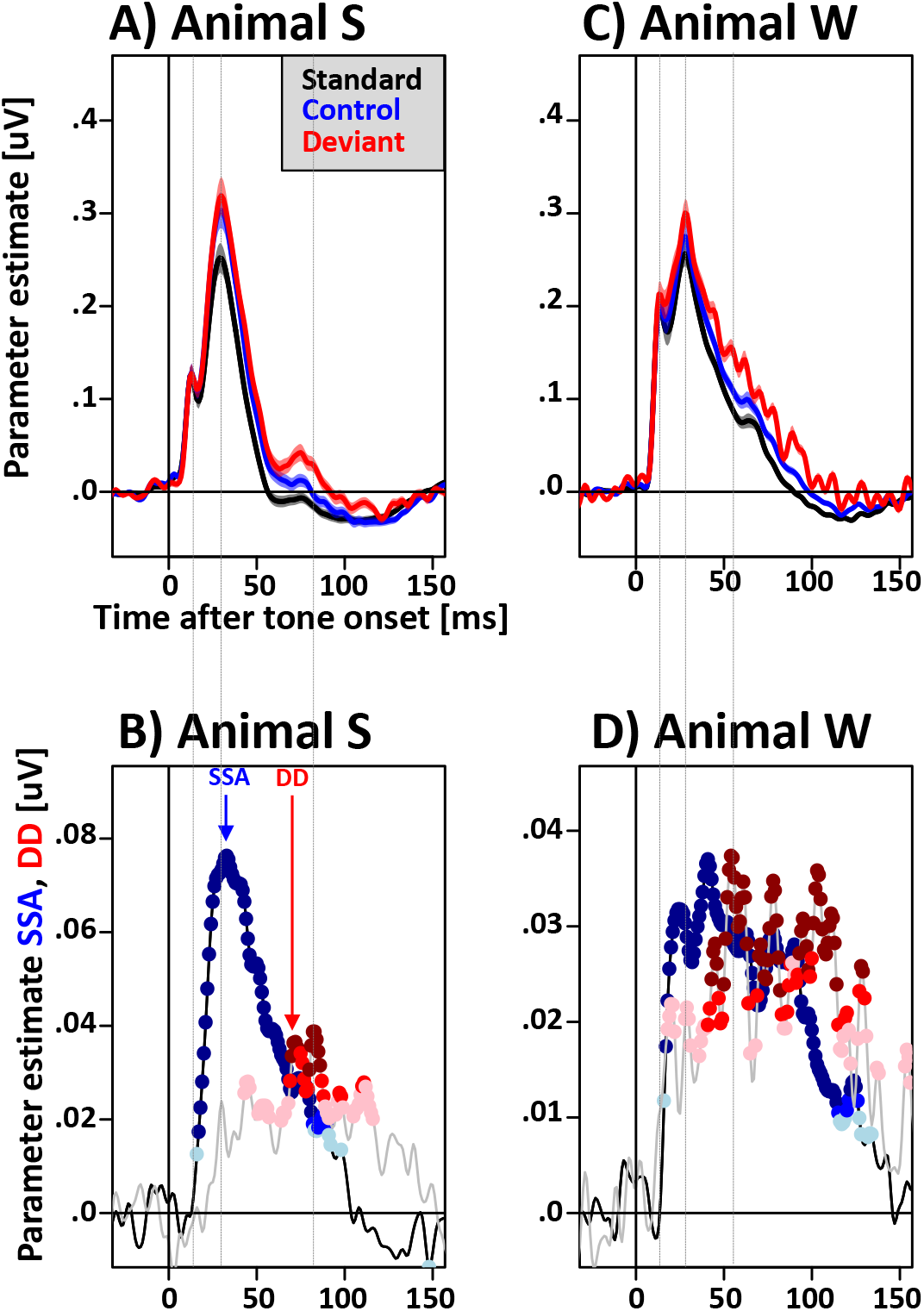
Multi-unit mismatch responses in the superior temporal plane. (A&C) Average multi-unit responses to the standard (black), the deviant (red) and the control tone (blue) as a function of time from tone onset. (B&D) SSA and DD as a function of time from tone onset. Blue and red dots of different shading indicate p-values of a time-resolved t-test.

Based on the multi-unit profiles, four epochs of interest were identified: activity of thalamic afferents in the time period from 8 to 12 ms after tone-onset (**th**), the first wave of evoked A1 activity from 25 to 45 ms (**evk**), a period of suppressed activity between 70 and 120 ms (**sup**), and a period of rebounding activity between 150 and 250 ms (**reb**). For each epoch of interest we also identified all contacts within specific layers of interest. For the epochs evk, sup and reb that included all contacts in putative layers 3,4 5 and 6. For epoch th we included all contacts below putative deep layer 3.

Figure 6 averages normalized MU activity in these time-windows for both animals separately. MU activity was normalized by regressing out all potential confounds including stimulus-onset asynchrony (*log(SOA)*), tonal frequency (*TF*), and the difference in tonal frequency (ΔTF) (see 4.9 and 4.10 for details). The results highlight the emergence of SSA and DD in the evoked period in both animals (Figure 6). Both signals are also present in the suppression period, but while SSA is weaker than in the evoked period, DD is stronger. Both SSA and DD were much weaker in the rebound period. However, DD remained significant for one of the two animals. For both animals, SSA also reached significance in the thalamic period, albeit with a rather small effect size. Figure 7A provides a more detailed look at the presence of the two signals using a finer temporal and spatial grid. This analysis shows that SSA can be identified in all putative layers 1 through 6. While SSA can be identified in all layers, p-values are lowest in layer 3, followed by layer 5, and layer 4. Layers 1, 2 and 6 only show comparatively week effect. Note that this layer-specificity is only partially driven by larger effect sizes. Figure 7B shows that layers 1,2 and 6 have smaller effect sizes. However, effect size is comparable in layers 3, 4 and 5. This suggest that differences in significance between these layers may be driven by higher variability in the prevalence of SSA responses in layer 5 and 4 MU populations. MU populations with significant DD response are restricted mostly to layers 3 and 4. Only in later parts of the response between 40 and 120 ms do DD responses emerge in layer 5 and 6. Neither SSA nor DD reach significance in the rebound time-window between 150 and 250 ms.

**Figure 6.**
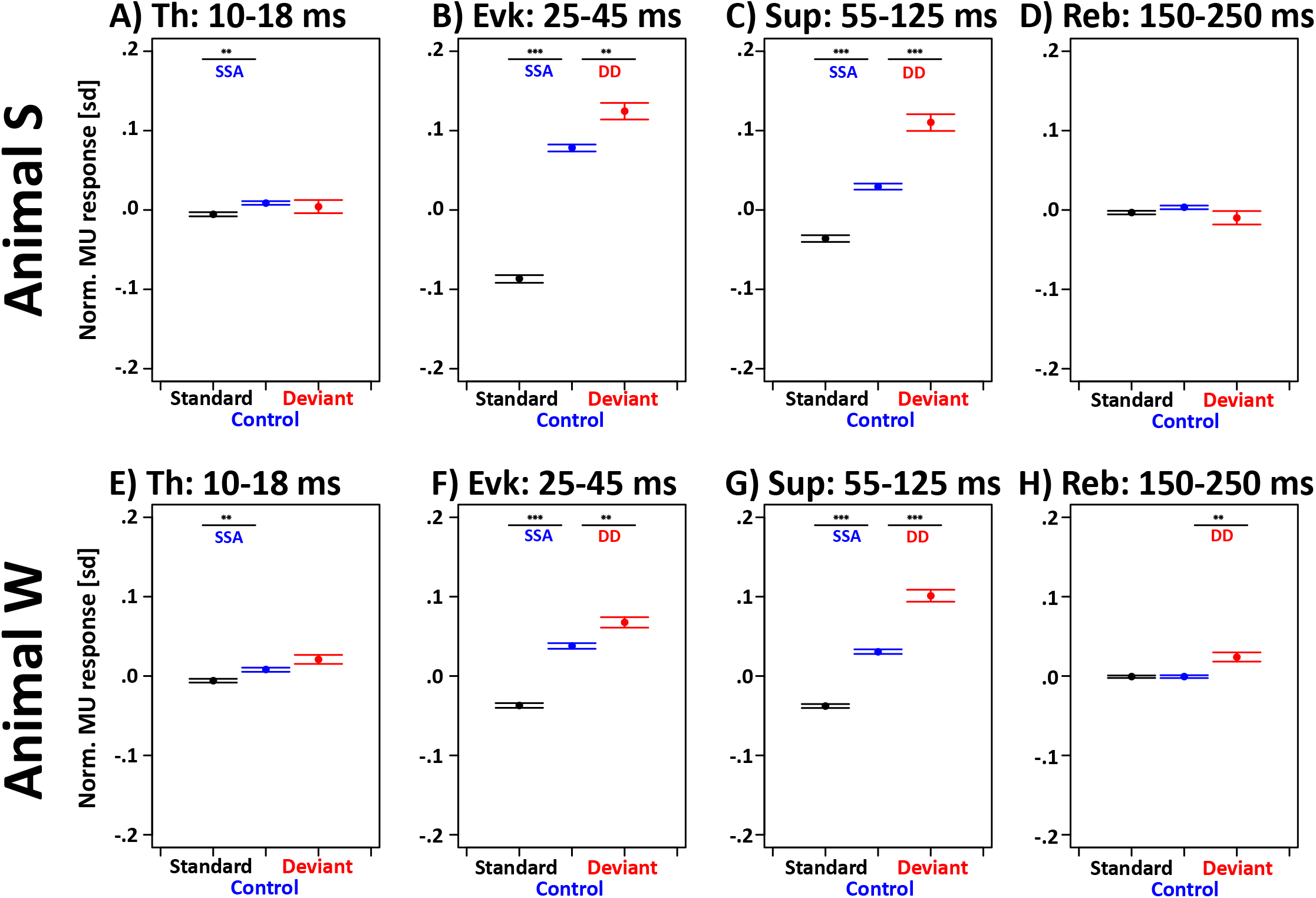
Average normalized multi-unit activity in response to standards, controls and deviants in four time-bins. Small but statistically significant SSA was observed in the thalamic period for both animals. SSA is largest in the evoked period, smaller in the suppression period and not significant in the rebound period. DD first reaches significance in the evoked period, and peaks in the suppression period.

**Figure 7.**
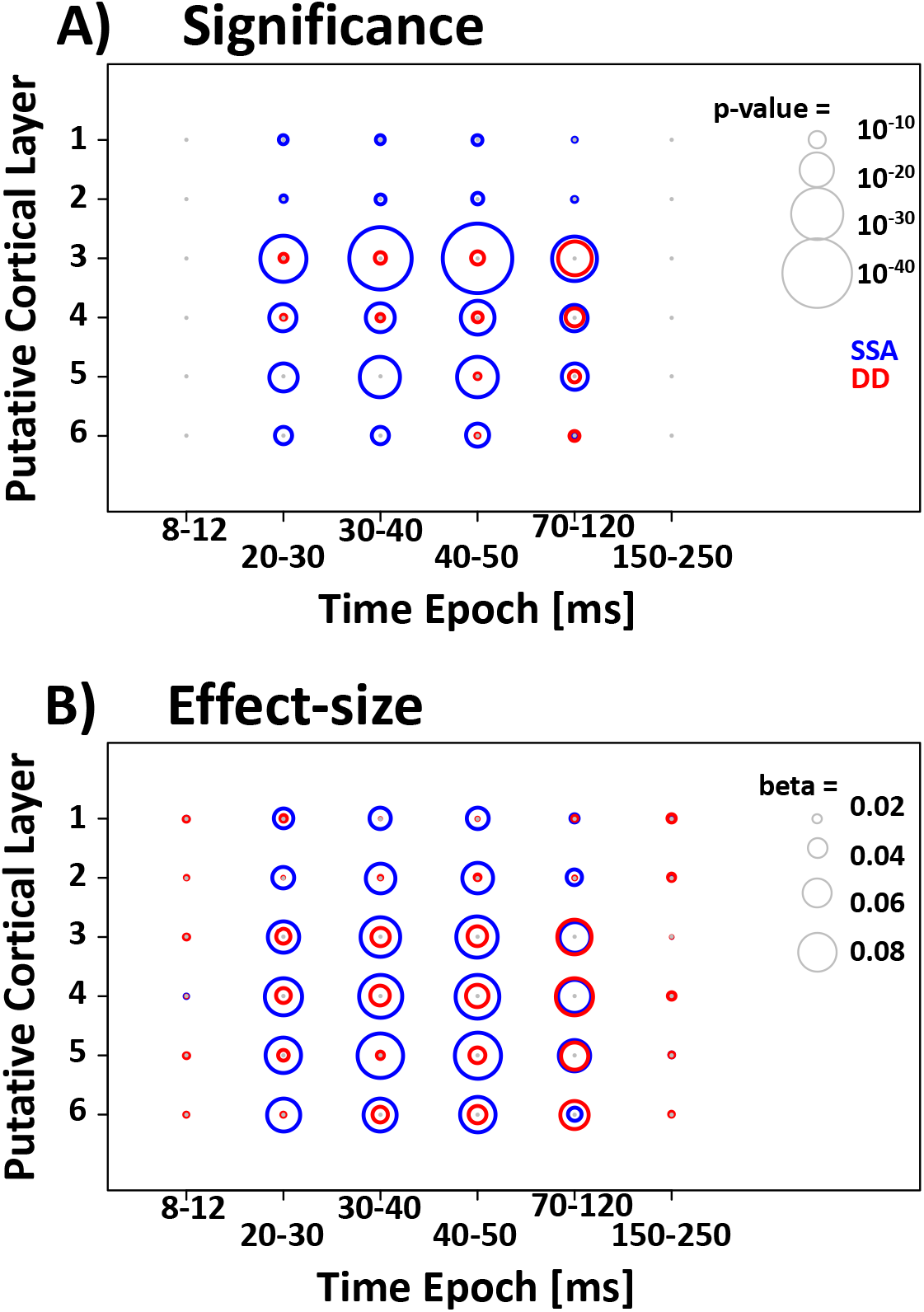
Significance and effect size of SSA and DD as a function of latency and putative cortical layer. (A) Color of the circles indicates the effect (blue=SSA, red=DD); size of the circles indicates the p-value of the tests. Size of the circles saturates for p-values below 10^−40^. (B) Size of the circles indicates effect size.

### 2.3 Adaptation model of monkey A1

There is a lively debate about whether experimentally measured DD is indeed dependent on active memory comparison processes or if it instead reflects a failure of the many-standard control condition to adequately correct for the level of SSA of the deviants. In particular, the early rising flank of DD which closely matches that of SSA (Figure 5) could point towards the later alternative. However, others have argued that the many-standard control condition may actually over-compensate for SSA, thus falsely concluding the absence of DD when actually it might have been present. The best way to resolve this concern is the use of a detailed computational model that accurately captures key aspects of adaptation in the data.

We thus set up a feed-forward model that uses short-term presynaptic depression (**STPSD**) at thalamo-cortical synapses to account for the observed multi-unit responses. The model was based on earlier work to account for adaptation in mouse primary auditory cortex (Nelken, 2014; Taaseh et al., 2011), and our own work to model refractoriness in auditory evoked EEG components in the monkey (Teichert et al., 2016). The model was set up to capture A1 receptive fields, as well as the three key properties of adaptation: (i) refractoriness, (ii) SSA, and (iii) their short-term habituation. Figure 8A depicts the architecture and the 5 parameters that determine the properties of the passive adaptation model. The parameter *σ_A1_* describes the range of thalamic cells from which a pyramidal cell receives it frequency-specific input. The parameter *a_Pedestal_* describes the relative contribution of stimulus-specific and non-specific input, and thus stimulus specific and non-specific adaptation. The parameters *U* and *τ* describe properties of short-term synaptic depression. *U* describes the cumulative vesicle release probability defined as the proportion of vesicles released by all the spikes elicited by a specific tone. This parameter determines properties of short-term habituation. The parameter τ describes the time-constant of vesicle recovery, and thus the rate at which adaptation dissipates. Lastly, the parameter *σ_Th_* describes the width of the thalamic receptive fields, and thus the band-width of the adaptation channel (see Methods for details).

**Figure 8.**
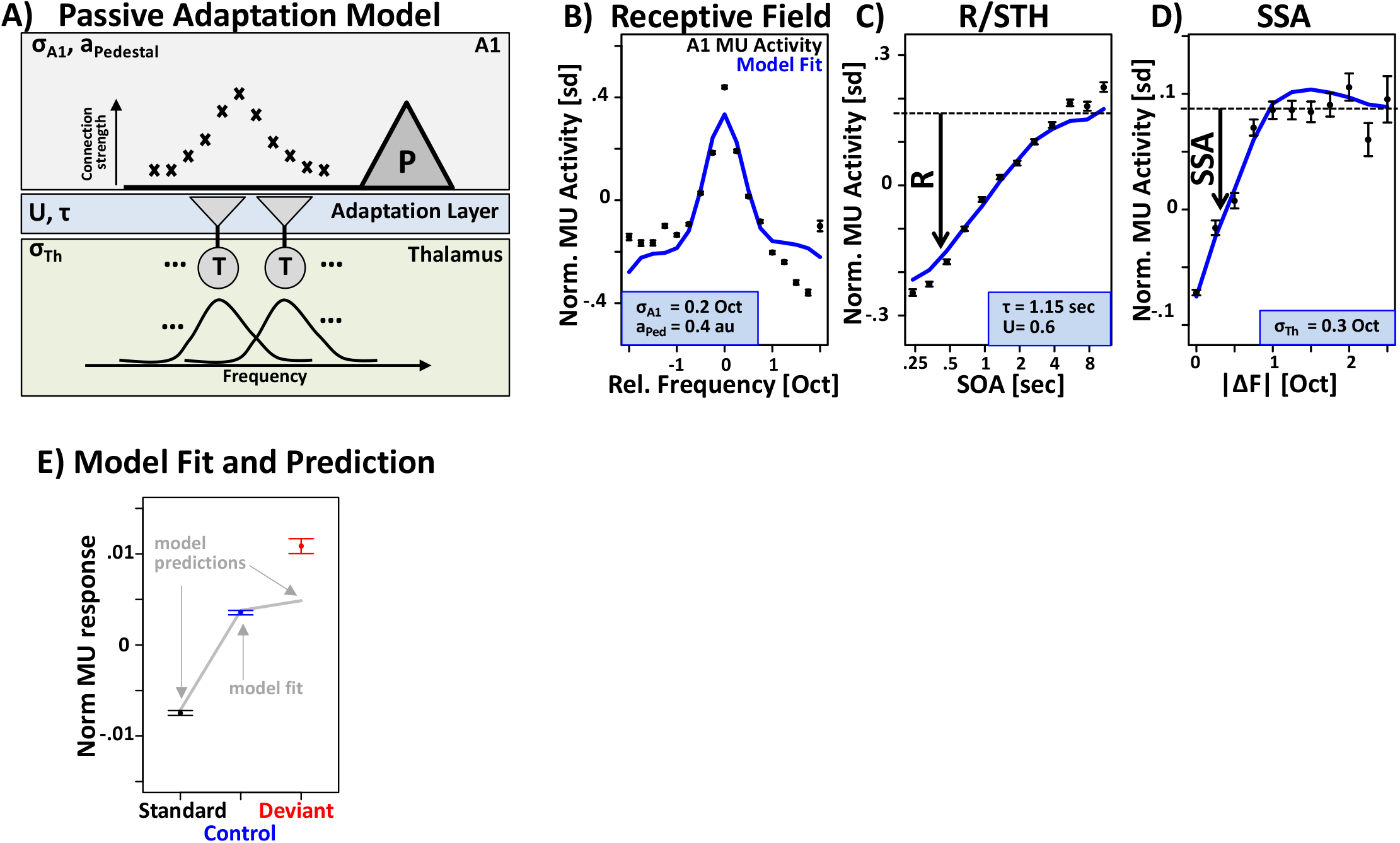
Adaptation Model. (A) The model assumes that pyramidal cells in A1 receive input from thalamic afferents via synapses that are subject to short-term synaptic depression with cumulative release probability U and recovery time constant τ. Additional parameters specify receptive field width of thalamic neurons, and the connection structure from thalamus to A1. The model was fit to the data from the many-standards control condition. (B) Multi-unit activity as a function of tonal frequency relative to preferred frequency of each multi-unit (black) and model fit (blue). (C) Multi-unit activity increases as a function of time since the last tone (black) and model fit (blue). (D) A1 multi-unit activity as a function of absolute tonal frequency difference between the past and current tone (black) and model fit (blue). (E) The recovered model parameters were then used to predict responses to standards and deviants to determine if adaptation can predict the actual data in these two conditions. Adaptation very closely predicted the attenuated response of the standards but failed to account for the enhanced responses of the deviants.

For each multi-unit, we extracted a tone-evoked response on a given trial by averaging activity in the time-window between 20 and 45 ms. Data from all trials and of all tone-responsive multi-units in both animals were z-scored and combined into one large data-frame. Because the model assumes a frequency-tuned receptive field, we only included multi-units that were significantly modulated by tonal frequency. This data set was then fit to the passive adaptation model using a modified gradient descent technique to identify the parameter values that minimized the residual sums of squares. Note that this fit included only data from the control condition, i.e., tones of unpredictable tonal frequency and onset time. Deviants and standards from the regular blocks were never presented to the model during the fitting process. Figure 8B depicts the average multi-unit receptive field (black) and the model fit (blue). Figure 8C depicts the time-course of recovery of multi-unit activity from refractoriness. Figure 8D depicts stimulus-specific adaptation of the multi-units as a function of similarity of subsequent tones, i.e., their frequency separation.

### 2.4 Release from adaptation cannot account for empirically observed DD

To test if adaptation in the control condition matches the degree of adaptation for the deviants, we simulated responses of the model in the regular condition that contains the deviants. Note that the model parameters were optimized by fitting them to the data in the control condition. Hence, responses of the model to the tones in the regular condition, i.e., the standards and deviants, are predictions, not fits. If the level of adaptation is matched between controls and deviants as intended by the many-standards control condition, then the model should predict identical responses to controls and deviants. Figure 8E (gray lines) shows that responses to the controls and deviants are indeed very similar. However, the results also show that responses to the deviants are slightly larger than to the controls, thus indicating that adaptation in the control condition was slightly weaker than adaptation for the deviants. However, the discrepancy of adaptation between the conditions is an order of magnitude smaller than the DD observed in the actual data depicted in red. Overall, this finding confirms the validity of the subtraction approach in combination with the many-standards control condition. Based on this validation, our findings provide strong support for the notion that the empirically observed DD is not just the result of a failure to adequately match adaptation of the controls to that of the deviants.

The results of our simulations validate the subtraction approach and suggests that two distinct mechanisms, adaptation and DD, contribute to the observed mismatch responses. This finding paves the way for a comparison of functional properties of the two mechanisms. The following sections thus provide a broader and more principled comparison of the functional properties of SSA and DD to test whether distinct functional properties confirm the notion that they indeed arise from distinct neural mechanisms. Ultimately, this may inform the specific roles of these two mechanisms for auditory short-term memory and/or pre-attentive guidance of attention.

### 2.5 SSA and DD have different time-course relative to stimulus onset

First, we conducted a principled comparison of the latency of all effects that may be mediated by STPSD in the context of our model, i.e., R, SSA and DD. If these components are indeed mediated by the same underlying mechanism there is a strong argument to be made that all three of them should emerge at the same latency in the actual data. We thus tested whether the time-courses of the parameter estimates for R, SSA and DD, i.e., *β_R_(t)*, *β_SSA_(t)*, and *β_DD_(t)* shared the same time-course or not. Figure 9A&D shows the time-courses of these three regressors as a function of time from stimulus onset for the two animals. Both *β_R_* and *β_SSA_* share a strikingly similar time-course that peaks at around 35 ms after stimulus onset and reaches a trough between 120 to 150 ms. Note that the two regressors themselves are uncorrelated. Hence, there is no a-priori reason that their parameter-estimates should share the same time-course. In contrast, the time-course of β_DD_ is clearly distinct with a peak at around 70 ms. The time-courses of all three parameter estimates were entered into a principal component analysis. In both animals, more than 99% of the variance was explained by the first two principal components. Figure 9B&E depict the time-courses of all three principal components. Note the similarity between the time-course of the first principal component and the time-courses of *β_R_* and *β_SSA_.* The second principal component has a delayed time-course much like *β_DD_*. Figure 9C&F plot the proportion of variance explained by each principal component for each of the three regressors. This analysis confirms that more then 95% of the variance of *β_R_* and *β_SSA_* can be accounted for by the first principal component. This same component also contributes between 20 and 50% of the variance to *β_DD_.* However, the remaining 50-80% of β_DD_ are accounted for by the second principal component. The contribution of this second principal component to *β_R_* and *β_SSA_* is negligible. The comparison of the time-courses thus supports the notion that R and SSA could be mediated by the same underlying mechanism, e.g., STPSD, that also contributes some, but not all of the observed effect of DD. In contrast, these results suggest that the same mechanism alone can not account for DD.

**Figure 9.**
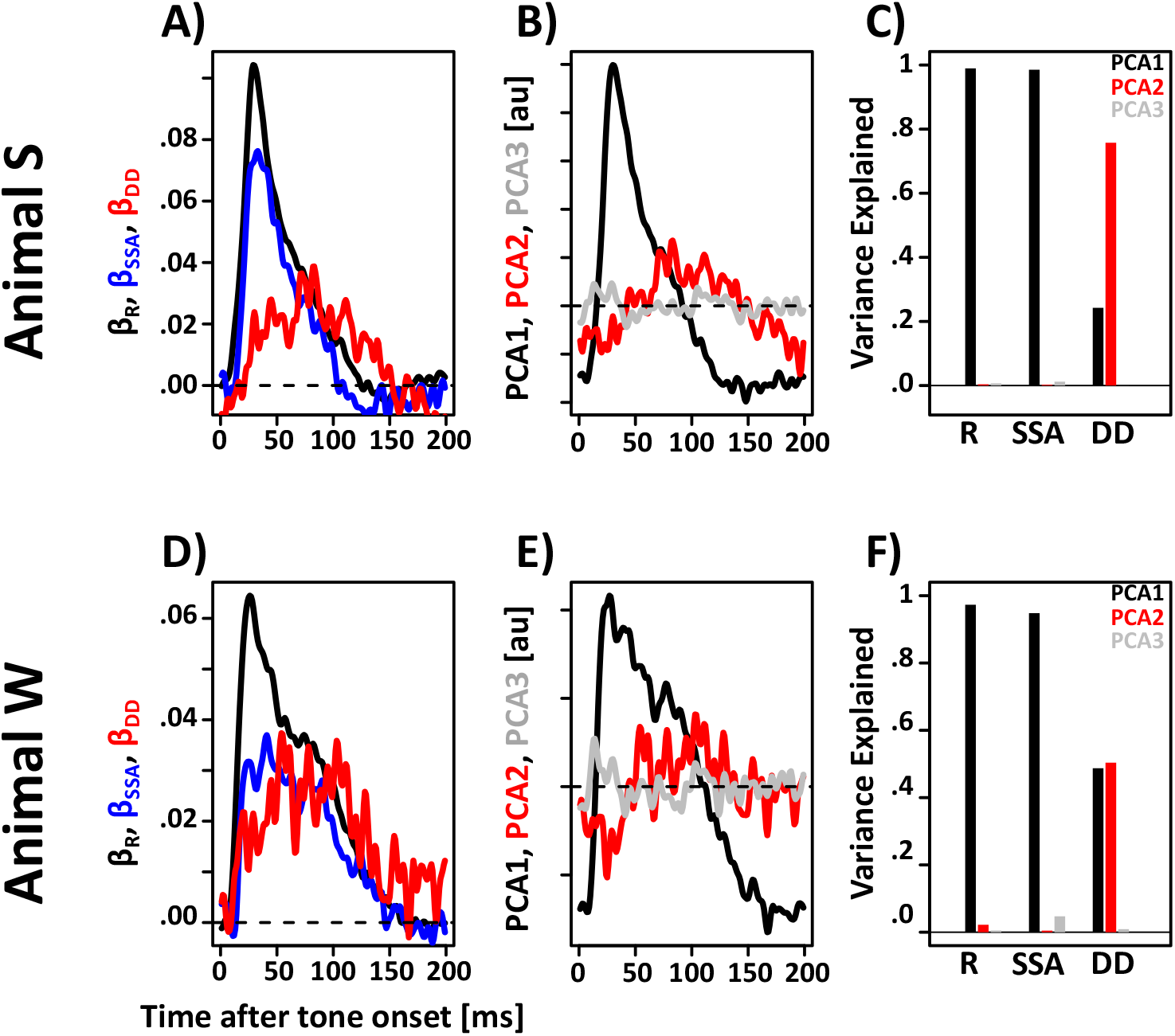
Comparison of latency of R, SSA and DD. (A&D) Time-resolved parameter estimates for R (black), SSA (blue) and DD (red). (B&E) Time-courses of the three components of a PCA analysis including the data from (A&D). (C&D) Proportion of variance explained by each of the three principal components. R and SSA are explained almost entirely by the first principal component. DD is explained by a mixture of the first and second principal component.

### 2.6 DD is stronger for larger frequency differences ΔF between standard and deviant

Next, we set out to determine how the magnitude of SSA and DD depends on the frequency difference ΔF between the standard and the deviant. If the two signals are mediated by distinct mechanisms it is possible that they have a different dependence on ΔF. Figure 10A-D shows responses to the standards, controls and deviants in the four response epochs. Note that in the evoked period responses to the control tones are mostly attenuated for |ΔF| below ~0.75 octaves. Responses to the deviants are similar to the controls for |ΔF| below ~0.75 octaves. However, responses for the deviants rise above the value of the controls mostly for values |ΔF| above 1 octave. A similar picture emerges for the suppression period. However, responses to the controls show more of a linear increase with |ΔF|. Figure 10E-H isolates SSA and DD using the subtraction approach. These subtractions confirm that SSA rises relatively briskly as a function of |ΔF| and then reaches more of a steady state or at least a rate of slower increase for |ΔF| > 0.75 octaves. In contrast, DD stays close to chance until |ΔF| reaches values of 1 octave or above. A similar pattern emerges for the suppression period. However, DD seems to start rising above chance for somewhat smaller values |ΔF|. Overall, these data suggest that SSA determines the bulk of MMN for values of |ΔF| below 1 octave. The additional increase of the MMN for larger |ΔF| values is carried by mostly by DD.

**Figure 10.**
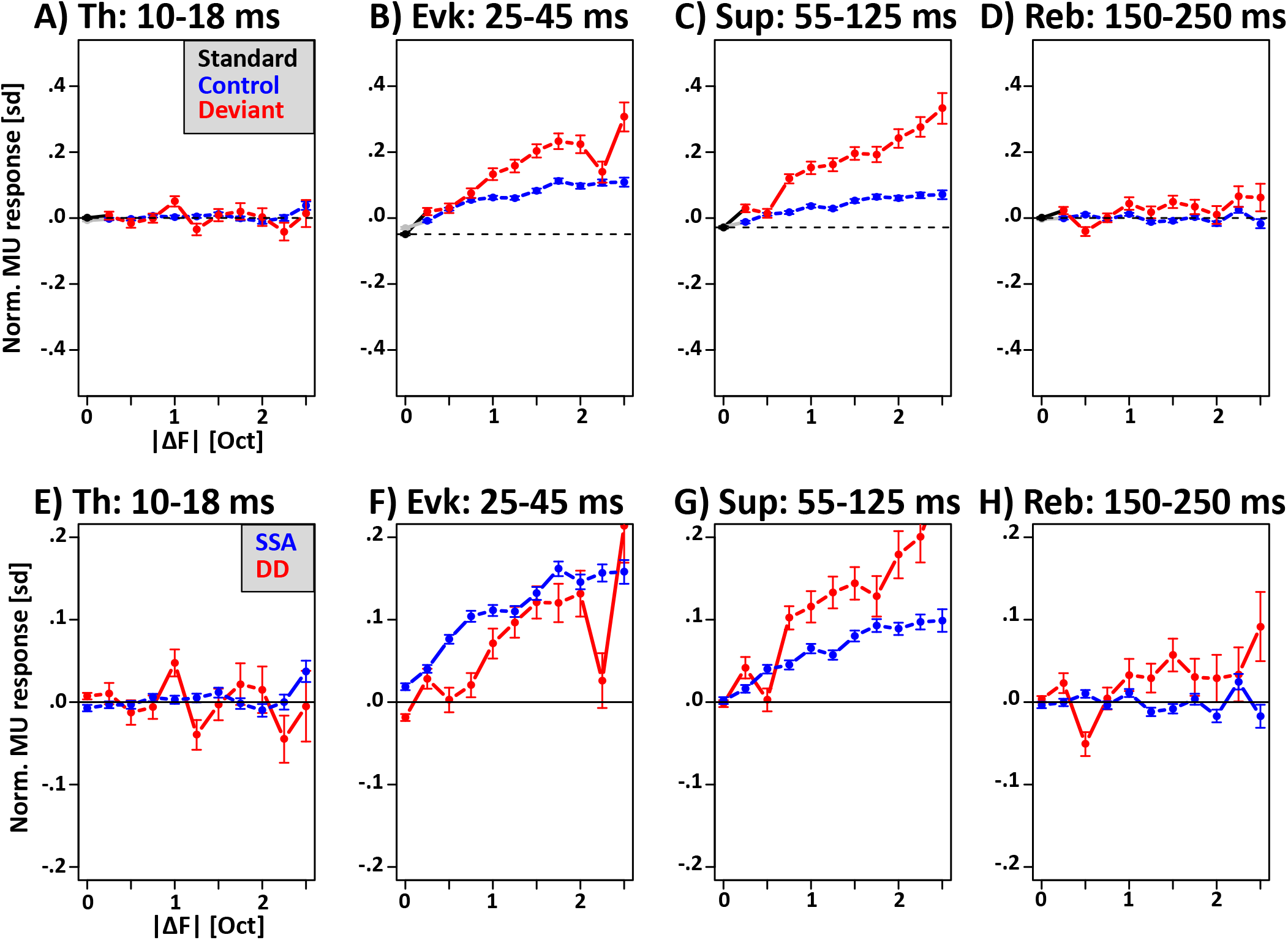
Comparison of frequency tuning of SSA and DD. (A-D) Normalized multi-unit response for standards, controls and deviants as a function of stimulus onset asynchrony in the four response epochs. Note that standards always have an |ΔF| equal to zero. (E-H) SSA, i.e., control – standard and DD, i.e., deviant – control as a function of |ΔF| in four response epochs. Note that SSA starts rising for smaller |ΔF| than DD in both the evoked and suppression period.

### 2.7 SSA and DD have different dependence on inter-stimulus interval

Next, we set out to compare another key functional property of R, SSA and DD, namely their dependence on the time between subsequent tones. If the three signals are indeed mediate by the same mechanism, e.g., STPSD, then they all should decay over time as synapses recover from depression. Figure 11A-D shows the average responses to deviants, controls and standards as a function of SOA in the four response periods. For all three stimulus types, activity increases as a function of SOA in the evoked and suppression period. Figure 11E-H plots SSA, i.e., the difference between controls and standards as well as DD, i.e., the difference between the deviants and the controls as a function of SOA. Two interesting observations emerge: First, the prevalence and magnitude of SSA is correlated with the prevalence and magnitude of R: Both, SSA and R, are small or completely absent in the thalamic and rebound period, intermediate in the suppression period and strongest in the evoked period. Second, in the evoked and suppression period SSA gradually diminishes for longer SOAs (Figure 11F&G), at about the same rate at which responses recover from refractoriness (Figure 11B&C). A different pattern is observed for DD. While refractoriness and SSA are strongest in the evoked period, DD is maximal in the suppression period. Most importantly, however, DD does not seem to depend on SOA as is the case for SSA. Taken together, these findings show that R and SSA decay over time at a comparable rate, while DD is not meaningfully affected by the delays used in our study (1/4 to 12 seconds).

**Figure 11.**
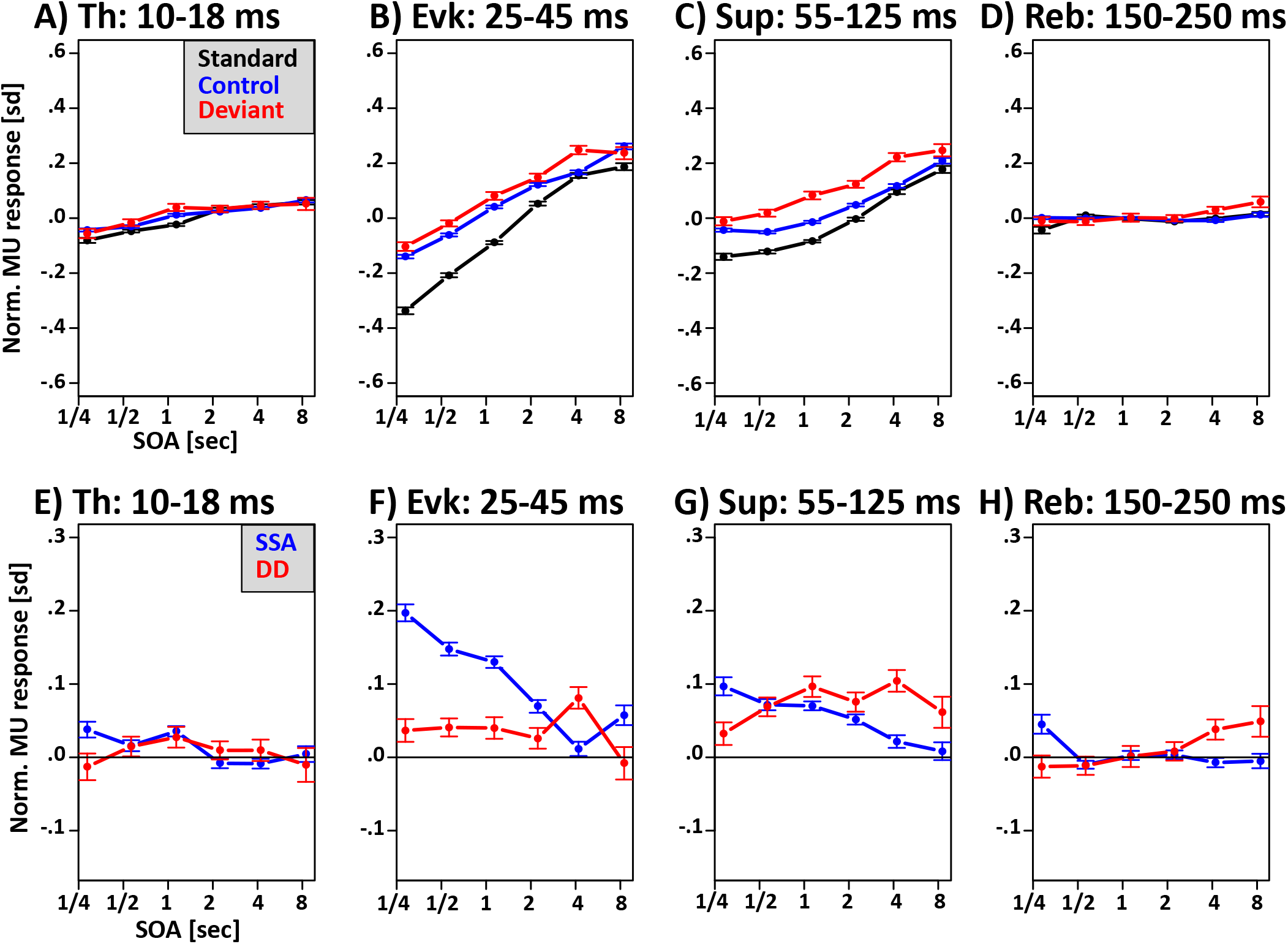
Comparison of recovery time-course of R, SSA and DD. (A-D) Normalized multi-unit response for standards, controls and deviants as a function of stimulus onset asynchrony in the four response epochs. (E-H) SSA, i.e., control – standard and DD, i.e., deviant – control as a function of stimulus onset asynchrony in four response epochs.

### 2.8 DD can be observed in multi- and single units that do not exhibit SSA

Lastly, we set out to analyze if R, SSA and DD preferentially affect the neural responses of the same or different units. The rational being that if DD is indeed the result of incomplete compensation for SSA, then all units that show DD would also have to show SSA. Figure 12A shows the percentage of MUs whose activity is significantly modulated by tonal frequency (**Frq**), R, SSA and DD in the four response periods. As expected for multi-units in primary auditory cortex, most were significantly modulated by Frq (gray bars). A considerable number of MUs (50%) were modulated by Frq even in the thalamic period. This is also expected given the known frequency specificity of thalamus. The lower fraction (relative to the evoked period) probably reflects the lower signal-to-noise ratio of the thalamic fibers, as well as the absence of fiber responses at all electrode contacts located above thalamic terminals in layer 4 and deep layer 3. In addition, the majority of MUs was also modulated by R reaching a peak of >90% in the evoked period. Note the relatively small amount of significant MUs in the thalamic period. This is consistent with the notion that R, at least when measured with recovery periods of >250 ms, is not inherited from thalamus, but rather generated in cortex or at thalamo-cortical synapses. A smaller fraction of MUs was modulated by SSA reaching a peak of 40% in the evoked period. DD was the least commonly observed effect peaking at around 15% in the suppression period.

**Figure 12.**
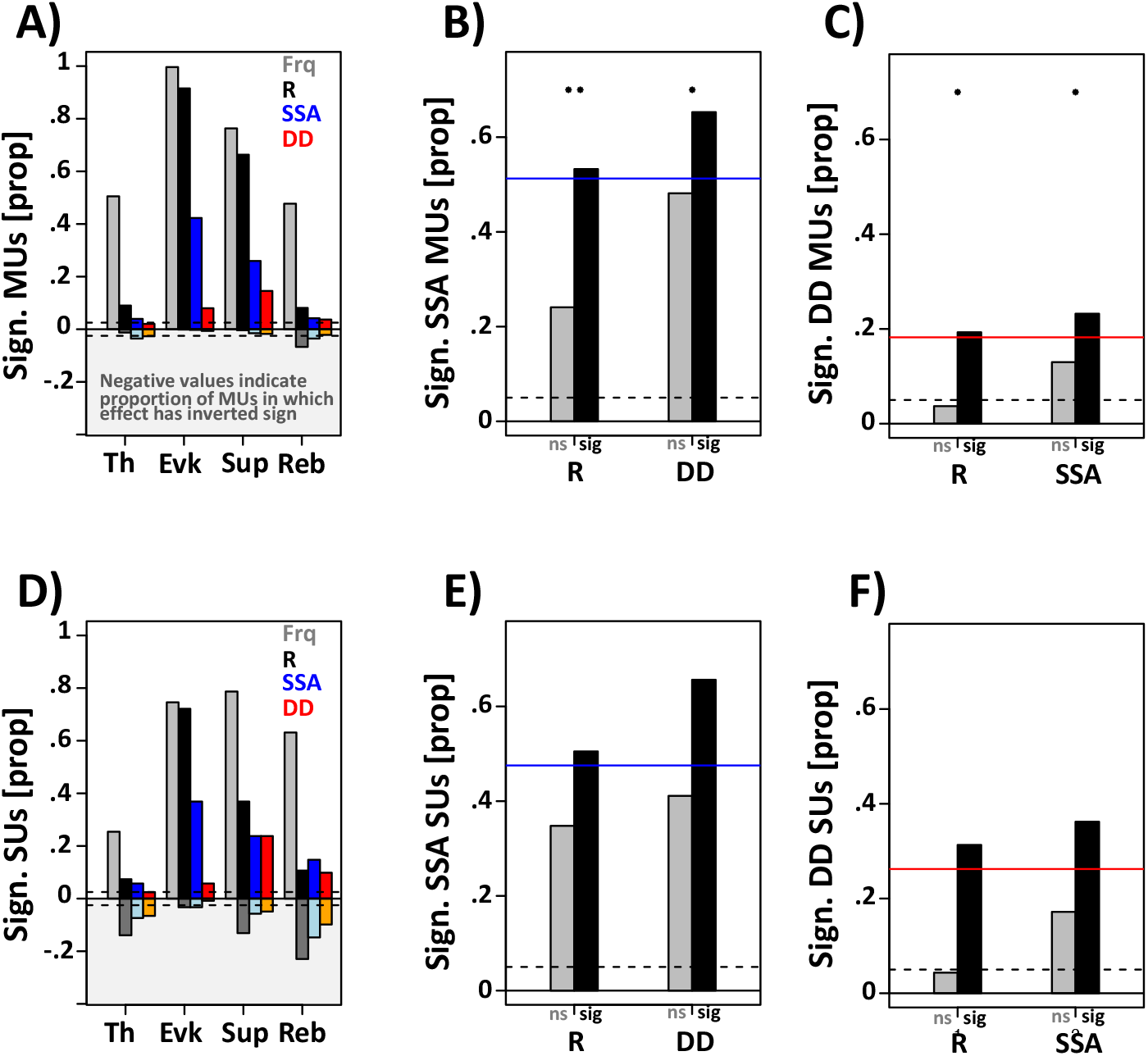
Multi- and single-unit response selectivity. (A) Proportion of multi-units significantly modulated by tonal frequency (Frq, gray), refractoriness (R, black), stimulus-specific adaptation (SSA, blue) and deviance detection (DD, red). Bars extending below zero indicate the proportion of cells with a significant effect of inverted sign. (B) Proportion of multi units exhibiting SSA given that the same multi-unit does (black) or does not (gray) exhibit R (left two bars) or DD (right two bars). Blue line indicates proportion of multi-units with SSA across the entire population. Asterisks above the bars indicate if the proportion of multi-units with SSA is significantly modulated by the presence of R or DD. (C) Same as (B) but for DD instead of SSA. (D-F) Same as (A-C) but for offline sorted single cells instead of multi-unit activity.

For subsequent analyses we defined a MU as modulated by R, SSA or DD if it was significantly modulated either in the evoked or the suppression period and the effect had the appropriate sign. Given the additional criterion of requiring the correct sign, there is a false positive rate of α=0.025 in each epoch.

Assuming independence of the two epochs, we end up with a conservative estimate of false positive rate of α=1-(1-0.025)^2=0.049. We round this value up to 0.05 to get an even more conservative estimate of the false positive rate for this compound test. Figure 12B shows the proportion of multi units with significant SSA depending on whether (black) or not (gray) it was significantly modulated by R or DD. Chi-square tests were used to determine whether or not the prevalence of SSA was modulated by the presence of the other two effects. The tests show that multi units are more likely to exhibit SSA if they are also modulated by R or DD. Similar results are shown for DD in Figure 12C. Two key findings stand out: (1) if a multi unit is not modulated by R, the chance of it exhibiting SSA or DD drops significantly; (2) an above-chance number of MUs are significantly modulated by DD even thought they are not significantly modulated by SSA. This analysis suggests that a significant fraction of multi units that are not affected by SSA, are nevertheless significantly modulated by DD.

To some extent, these findings may reflect the fact that a multi unit can reflect responses of many single units with distinct response properties. To test whether the results hold up for single cells, we repeated the analyses using offline sorted single cells. Figure 12D-F shows qualitatively identical and quantitatively very similar result for offline-sorted single cells. These results confirm the notion that a cell can exhibit DD even if it does not exhibit SSA. It should again be noted that in some of the cells that exhibit DD but not SSA, the lack of SSA may be due to a false negative.

### 2.9 Recording locations on superior temporal plane

We finally explored whether R, SSA and D were distributed differentially across the sampled surface of the superior temporal plane. Figure 13A&D shows the best frequency at the estimated location of each penetration. Frequency preference was measured with a standard tonal response field paradigm. This data set also includes recording sessions that were not used for this experiment. It thus covers a larger chunk of auditory cortex and more recording sessions. In animal S, the tonotopy shows a clear gradient from high to low frequencies as the recording locations progress from posterior to anterior. There is also an indication of an inverted gradient for even more anterior recording locations. The direction of the main gradient is consistent with the notion that most recordings were located in primary auditory cortex. For animal W a tonotopic gradient is harder to discern. Nevertheless, the few recording locations with the highest preferred frequencies are relatively far posterior. Hence, it is likely that the recording locations were also mostly in primary auditory cortex. Figure 13B&E show the location of multi-units that were/ were not significantly modulated by R (black/ gray), SSA (blue/ gray) or DD (red/ gray). Overall, all of the effects seem to be present across the entire sampled surface of the superior temporal plane. In animal S the two most posterior recording locations seem to be least modulated by both SSA and DD. However, more dense sampling would be necessary to establish a significant antero-posterior gradient of SSA and DD.

**Figure 13.**
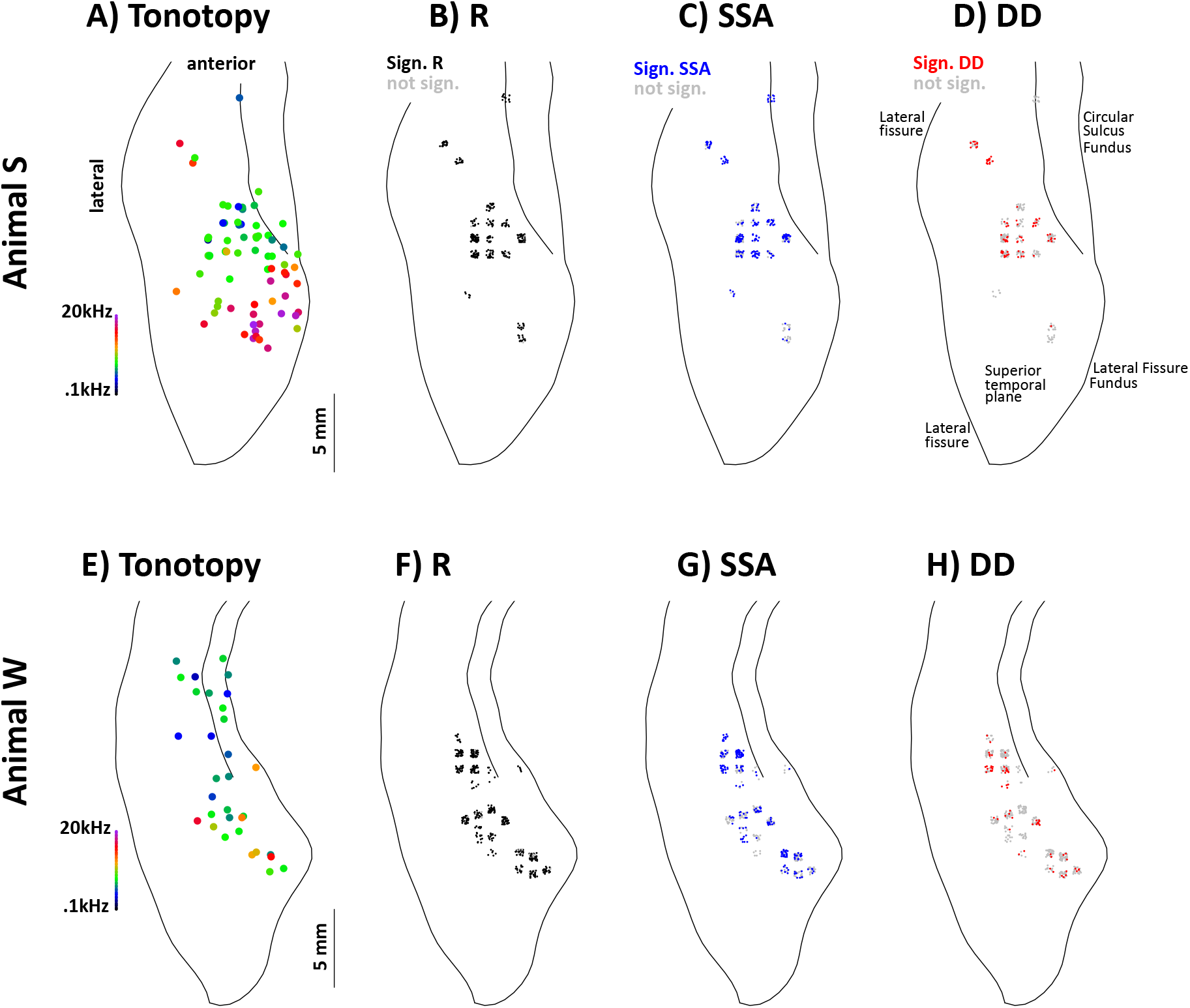
Topography of R, SSA and DD. (**A**&**E**)> Approximate location and preferred frequency of all recording locations in the superior temporal plane. (B&F) Location of multi-units that are either modulated by R (black) or not (gray). (C&G) Location of multi-units that are either modulated by SSA (blue) or not (gray). (D&H) Location of multi-units that are either modulated by DD (black) or not (gray).

## 3 Discussion

There is an ongoing debate about whether MMN can be accounted for entirely by passive adaptation, active memory-comparison processes/ predictive coding such as deviance detection, or a mix of both. Here we provide evidence that micro- as well as macroscopic mismatch responses in the rhesus are dominated by adaptation at short latencies but also provide support for DD, a putative active memory-comparison mechanism, that emerges at longer latencies. Most importantly, we show that adaptation and DD have complementary functional profiles in line with distinct yet complementary roles in mediating auditory short-term memory and the prioritization of informative auditory events. In the following, we will discuss the implications of these key findings in more detail.

### 3.1 Micro- and Macroscopic deviance detection in the monkey

To our knowledge, the data presented here is the first direct evidence of DD in the macaque. The impact of this finding is further amplified by the fact that DD was found at the macro- and microscopic level, and that the timing at both levels showed the same temporal progression from early SSA to later DD that has been observed in the human. At the macroscopic level, DD was detected as an increase of the P55, a surface positive mandatory EEG component that peaks over fronto-central electrodes around 55 ms after stimulus onset. SSA could be detected at shorter latencies as a reduction of the P21 and P31, two surface positive mandatory EEG components that also peak over fronto-central electrodes around 21 and 31 ms after stimulus onset. At the microscopic level, DD was identified in A1 multi-unit activity mostly in layer 3, 4 and 5. DD peaked around 70 ms after tone-onset. SSA could be observed much earlier, reaching peak amplitudes around 30 ms after tone-onset. Finally, DD was also identified in offline sorted single units.

Recent work by Fishman and Steinschneider have used similar experimental paradigms but did not find evidence for DD in monkey primary auditory cortex (Fishman, 2014; Fishman and Steinschneider, 2012). The following outlines several potential explanations for the discrepant results. (1) <u>Absolute tonal frequency</u> <u>differences</u> |ΔF|. In our hands DD started to emerge only if |ΔF| was larger than ~1 octave. Fishman and Steinschneider used a dynamic procedure that adjusted |ΔF| individually for each recording location to elicit about half the response of the best frequency. This value was likely below 1 octave that was necessary to robustly drive DD in our hands. (2) <u>Stimulus onset asynchrony</u>. In our hands MMRs were dominated by SSA at short SOAs. Hence, the use of 658 ms long SOAs by Fishman and Steinschneider may have led to a relatively small contribution of DD to the total MMR, thus making it harder to detect. (3) <u>Auditory Paradigm.</u> Our study used a variant of the roving standard paradigm in which each deviant becomes the new standard, thus dividing blocks of the regular condition into short runs of ~5-15 repetitions of identical tones of the same tonal frequency. Furthermore, the different runs were not only marked by different tonal frequencies, but also by different presentation rates, which may have further strengthened the perceptual segregation into short runs. Thus, the DD identified in our study may either reflect the detection of the deviant, as implied by the term DD, or it may mark the onset of a perceptually segregated new run. In contrast, Fishman and colleagues used the classic oddball and flip-flop paradigm. Neither of these paradigms lend themselves to being perceptually segregated into different runs.

A more recent study by Lakatos and colleagues reported MMR in the auditory cortex of the macaque using an oddball paradigm without a control condition (Lakatos et al., 2019). This prevented a direct dissection of the MMR into SSA and DD as was performed here and in the work by Fishman. However, multi-unit MMR by Lakatos and colleagues likely reflected at least in part DD because it could also be observed when both deviant and standard were far from the best frequency of the recording location, a condition that failed to elicit significant repetition suppression at the population level. Hence, at the level of multi-unit activity the two studies seem to come to convergent results. However, an interesting dissociation emerges at the level of frequency tuning of the putative DD component. In our hands, the bulk of DD only emerged when |ΔF| was larger than 1 octave. Lakatos and colleagues saw their effects at a fixed |ΔF| of 3 semi-tones, i.e., one quarter of an octave. A second interesting dissociation emerges at the level of current source densities. Lakatos and colleagues show a reduction of amplitude or even inversion of polarity of the supra-granular generators for the deviants compared to the standards. In our hands, to opposite seems to be true: responses of the supra-granular generators are stronger for the deviants compared to the standards (data not shown), a finding that is consistent with previous studies (Fishman, 2014; Fishman and Steinschneider, 2012; Javitt et al., 1996). Furthermore, while less consistent across animals, our data suggest that part of that increase in supra-granular generator amplitude can be attributed to DD.

### 3.2 Complementary functional profiles of mismatch responses mediated by adaptation and DD

Several of our analyses identified complementary functional profiles for mismatch responses mediated by adaptation and DD, respectively. The two adaptive components, R and SSA, emerged at short latencies and largely overlapped with the first wave of evoked cortical multi-unit activity. In contrast, DD emerged at longer latencies and largely overlapped with the subsequent period of suppressed cortical multi-unit activity. Both R and SSA dissipated almost completely if the time between subsequent tones was larger than ~4 seconds. In contrast, DD seemed to be largely invariant across the range intervals tested here. Finally, the effect of adaptation as a function of the absolute frequency difference |ΔF| between deviant and standard began leveling off at one octave. In contrast, DD mostly boosted responses if |ΔF| was larger than one octave. In summary, these results suggest that the contribution of adaptation to mismatch responses is strongest at short latencies, for short inter-stimulus intervals, and when deviants and standards are physically similar. The opposite is true for DD. Our findings complement recent work by Lakatos and colleagues which can be interpreted in a way that DD is broadcast across the entire tonotopic map of A1, while SSA remains localized to a relatively narrow range of the tonotopic map(Lakatos et al., 2019).

Relatively stronger neural responses to deviants have been implicated in the prioritization of potentially relevant sounds (Jääskeläinen et al., 2004). Hence, the complementary functional profiles of adaptation and DD suggest that the joint action of both mechanisms would facilitate a more versatile detection and prioritization of potentially informative auditory events. Release from adaptation would enable the tagging of potentially relevant information with very limited computational overhead, at very short latencies, and even for relatively similar tones. However, the temporal scope of this passive mechanism would be limited to relatively short time-scales below ~4 seconds, and it would fail to prioritize potentially informative events such as unexpected stimulus repetitions. In contrast, DD would enable integration of information over longer time-scales and tag potentially informative stimuli independent of their level of adaptation. However, this flexibility might come at a cost of more computational overhead, longer latencies and reduced stimulus sensitivity.

Similarly, the MMN has been intimately linked with auditory short-term memory (Näätänen et al., 2005). Our recent work on auditory short-term memory in the rhesus macaque has identified two functionally distinct memory components: a short-lived component that supports relatively high discriminability which may be homolog to echoic memory in the human, and a longer-lived component with lower discriminability that improves with repeated presentation of the standard which we have tentatively referred to as auditory recognition memory (Teichert and Gurnsey, 2019). We speculated that adaptation mediates echoic memory by leaving a short-lived ‘negative’ trace of past sounds mostly in primary auditory cortex, potentially in the form of depleted vesicles at synapses of neurons that fired to the previous sound. Furthermore, we speculated that recognition memory may be mediated in higher auditory regions such as rostral superior temporal gyrus and the ventral prefrontal cortex. Based on the similarity of their respective functional profiles, it is tempting to associate echoic memory with adaptation and auditory recognition memory with DD.

This interpretation provides an interesting take on a valid criticism of the memory account of the MMN that has been brought forth by May and Tiitinen (May and Tiitinen, 2010). They have argued that every sound is expected to leave a memory trace, even if it is presented just once. Hence, if MMN is indeed a memory-comparison process, it should be present regardless of how often a tone has been repeated prior to the presentation of a deviant. This, of course is at odds with the finding that the MMN only emerges if the standards are presented repeatedly. In line with the premise of May and Tiitinen, we argue that in the monkey, the MMN is consistent with the notion of reflecting not just one but two memory system. Adaptation is a neural correlate of (echoic) memory that indeed emerges after the very first presentation of a tone. In contrast, DD is a correlate of recognition memory, which strengthens with repeated presentation of a tone and reaches its maximum only after many repetitions. This gradual strengthening of recognition memory may explain why DD is only observed if the standard is repeated multiple times.

### 3.3 New evidence for DD

Our conclusion that the MMN is comprised of two mechanisms, i.e., adaptation and DD, is not novel per se (Jacobsen et al., 2003; Jacobsen and Schröger, 2001; Lakatos et al., 2019; Maess et al., 2007; Opitz et al., 2002; Schröger and Wolff, 1996). However, currently available evidence in support of the contribution of an active memory-comparison process, i.e., DD, has been challenged on various grounds. The work presented here strengthens the argument in favor of DD along three main arguments. (1) It provides a conclusive quantitative analysis that the longer latency of the MMN in general, and DD in particular, cannot simply be accounted for by stronger and longer-lasting responses from less refractory elements of the auditory system; It provides a quantitative argument that the control condition does a good enough job at matching adaptation of the deviants and thus confirming the validity of the subtraction approach; (3), it shows critical functional distinctions for the two extracted components thus further supporting the case for two separate mechanisms.

1. A key criticism of the memory account is that the MMN may not be a separate process, but rather the difference of two N1 responses whose different magnitude and duration reflect different levels of refractoriness of different parts of auditory cortex. Because studies so far have never explicitly incorporated the effects of refractoriness into studies of MMN, this criticism has been hard to refute quantitatively. To address this concern, we directly measured the effect of different levels of refractoriness on the magnitude and duration of various tone-evoked neural responses. As expected, refractoriness had a strong effect on multi-unit responses in auditory cortex. However, the effect of refractoriness was mostly limited to relatively short latencies. The presumed DD component emerged at longer latencies at which the effect of refractoriness was already waning. Furthermore, our data show that this is not only true for non-specific adaptation, i.e., refractoriness, but also stimulus specific adaptation, i.e., SSA. Together, this provides new and strong evidence that the longer latency of DD cannot be accounted for by stronger and longer responses by less adapted elements of the auditory system.
2. A second key concern is that adaptation for the control tones may be stronger than adaptation for the deviants, thus invalidating the key assumption of the subtraction approach. To address this concern, we developed a computational model that captured key aspects of adaptation in auditory cortex. The model was fit to the multi-unit responses in the control condition to determine (i) the tuning width of the stimulus-specific adaptation channel, (ii) the relative contribution of stimulus-specific to non-specific adaptation, (iii) whether the effects of subsequent tones sum linearly or sub-linearly, and (iv) how quickly adaptation dissipates during periods of silence. To our understanding, this is the most detailed model of adaptation that has explicitly been fit to auditory cortex responses using a gradient descent technique. To the extent that it accurately captures key aspects of adaptation in auditory cortex, it was possible to ask the model whether or not adaptation in the control condition matched the level of adaptation for the deviants. This effort revealed two main conclusions. First, it served as a reminder that this concern is very valid and cannot easily be addressed by qualitative arguments alone, i.e., without a sophisticated computation framework. Second, it revealed that while the level of adaptation was not perfectly matched between conditions, the difference was an order of magnitude smaller than the empirically observed DD.
3. A third concern is whether or not the two putative components, ie., adaptation and DD, indeed show distinct functional properties as would be expected for two different mechanisms. Differences between SSA and DD with respect to timing and topography have been identified previously but are relatively small. To strengthen this argument, we quantified other aspects of the two putative mechanisms. As outlined above, our results support the notion of different timing and additionally suggest different frequency tuning and life-time for the two putative mechanisms. Taken together, this strengthens the notion that the subtraction approach was able to pull out the contribution of two separate mechanisms as indicated by a number of genuinely distinct functional properties.

### 3.4 EEG Mismatch Positivity

This study is the first to report surface-positive mismatch responses in monkey EEG. Earlier studies in the monkey have not reported this component. Nevertheless, the finding is not completely unexpected. (1) Closer inspection of the mismatch responses in an earlier study (Honing et al., 2012) reveal a similar mismatch positivity around the time of the monkey P1 which corresponds to our P21 and P31. This early mismatch positivity was not included in the statistical analysis of the earlier paper so it is unclear if it would have reached significance. (2) Our previous work has shown that the early positive EEG components (P21, P31 and P55) are strongly and significantly affected by R. We had speculated that this non-specific reduction in response magnitude would also include a stimulus-specific component, thus giving rise to a mismatch positivity driven by SSA. (3) Multi-unit activity and current sources in monkey primary auditory cortex have provided ample evidence for SSA at relatively short latencies peaking around 30 ms after tone onset (Fishman, 2014; Fishman and Steinschneider, 2012; Javitt et al., 1994). Given the close relationship between the auditory cortex activity and fronto-central EEG responses, especially at short latencies, it would have been rather surprising not to identify such an early mismatch positivity. (4) The identification of a mismatch response that emerges at shorter latencies than the classical MMN is also expected from recent findings of mismatch responses in humans that arise earlier than the classical MMN around the time of the Nb, a midlatency component peaking around 40 ms after tone onset (Alho et al., 2012; Grimm et al., 2011; Grimm and Escera, 2012).

## 4 Material and Methods

### 4.1 Subjects

The EEG experiments were performed on 4 adult male macaque monkeys (macaca mulatta, animals R, W, J and S). The laminar recordings in A1 were performed in a subset of two of these animals (animals W and S). The treatment of the monkeys was in accordance with the guidelines set by the U.S. Department of Health and Human Services (National Institutes of Health) for the care and use of laboratory animals. All methods were approved by the Institutional Animal Care and Use Committee at the University of Pittsburgh. One animal (animal R, 10.5 kg, 12 years) had previously performed simple cognitive tasks in a different lab. Preliminary data collected from animal R helped optimize the EEG setup and auditory paradigms used in this study. The other three animals were between 5 and 6 years old and weighed between 8-9 kg at the time of the EEG experiments. Up until the first day of recording for the EEG experiments, these animals were not exposed to any of the auditory paradigms. After the EEG recordings, intracranial recordings were performed in animals S and W over a period of ~3 years.

### 4.2 Cranial EEG recordings

Details of the cranial EEG recordings have been reported previously(Teichert, 2016; Teichert et al., 2016). Briefly, EEG electrodes manufactured in-house from either medical grade stainless steel (monkeys J and W) or medical grade titanium (monkeys R and S) were implanted in 1mm deep, non-penetrating holes in the cranium. All electrodes were connected to a 32-channel Omnetics connector embedded in acrylic at the back of the skull. The different animals had between 21 and 33 electrodes implanted that formed regularly-spaced grids covering roughly the same anatomy covered by the international 10-20 system. Some of the EEG data analyzed here has been used in previous manuscripts(Teichert et al., 2016). There is no overlap between the analyses performed in the two previous manuscripts.

### 4.3 Laminar recordings in primary auditory cortex

Neural activity was recorded with 24 and 32 channel laminar electrodes (V-Probe, S-Probe from Plexon) positioned approximately perpendicular to the orientation of the left superior temporal plane. Early on, electrodes were inserted through 3mm diameter burr-holes within the recording chamber. Later on, the entire cranium within the recording chamber was removed. Recordings were referenced to the most posterior EEG electrode on the midline (Oz).

### 4.4 Experimental Setup

All experiments were performed in two small (4’ wide by 4’ deep by 8’ high) sound-attenuating and electrically insulated recording booths (Eckel Noise Control Technology). Animals were positioned and head-fixed in custom-made primate chairs (Scientific Design). Neural signals were recorded with a 256-channel digital amplifier system (*RHD2000, Intan)*. For the EEG recordings, neural signals were recorded at a sampling rate of 5 kHz. For the intracranial recordings, neural signals were recorded at a sampling rate of 30 kHz.

Experimental control was handled by a windows PC running an in-house modified version of the Matlab software-package *monkeylogic*. Sound files were generated prior to the experiments and presented by a sub-routine of the Matlab package *Psychtoolbox*. The sound-files were presented using the right audio-channel of a high-definition stereo PCI sound card (M-192 from M-Audiophile) operating at a sampling rate of 192 kHz and 24 bit resolution. The analog audio-signal was then amplified by a 300 Watt amplifier (QSC GX3). The amplified electric signals were converted to sound waves using either two insert earphones with foam plugs (ER2, Etymotic) (monkeys J and R), or a single element 4 inch full-range driver (Tang Band W4-1879) located 8 inches in front of the animal (monkeys W and S).

To determine sound onset with high accuracy, a trigger signal was routed through the unused left audio channel of the sound card directly to one of the analog inputs of the recording system. The trigger pulse was stored in the same stereo sound-file and was presented using the same function call. Hence, any delay in the presentation of the tone also leads to an identical delay in the presentation of the trigger. Thus, sound onset could be determined at a level of accuracy that was limited only by the sampling frequency of the recording device (5kHz: corresponding to 0.2 msec; 30kHz: corresponding to 33 μsec).

### 4.5 Auditory Paradigm of EEG study

Details of the auditory paradigm for the EEG recordings have been reported previously(Teichert, 2016; Teichert et al., 2016). Briefly, the study used 80 dB, 55 ms pure tones with a 5ms linear rise/fall times. During each experimental session, we presented 11 different frequencies spaced linearly in log_2_-space. The lowest tone was randomly selected from the interval between 500 and 800 Hz; the highest tone was drawn randomly from the interval of 3000 to 4000 Hz. Time between individual tone onsets (stimulus-onset interval, **SOA**) ranged between 0.2 and 12.8 seconds. The actual SOAs were drawn from a mix of two truncated exponential distributions. The time-constants of the two distributions were randomly selected from the interval between 300 and 600ms, as well as 1000 and 2000 ms, respectively. Tone presentations were structured into blocks between 9 and 12 minutes duration (**Fig 3B**). A block was terminated after the randomly determined time-criterion or after 600 tone-presentations, whichever happened first. The program generated two sequences of random numbers that determined SOA and tone identity (a number between 1 and 11 that corresponded to one of the 11 previously generated sound-files). A sequence of binary values determined whether the specific combination of tonal frequency and SOA that determined the last tone would be repeated for the next tone, or whether the next tone would be determined by the next pair of SOA and tone identity. In the so-called ‘random environment’, the likelihood of repeating a tone identity/SOA combination was 10%. In the ‘regular environment’, the likelihood of repetition was 90%, and the time as well as identity of the upcoming tone could be predicted with high accuracy. Blocks of random and regular environment were presented in an alternating order.

### 4.6 Auditory Paradigm of intracranial study

Key design choices for the auditory paradigm in the intracranial recordings were made based on the notion that intracranial recordings would be most informative to the EEG findings if they used the same or at least very similar experimental paradigm. Consequently, changes to the experimental design were kept to a minimum and were implemented only if required by the different recording situation. (1) To account for the limited number of trials that can be acquired with acute electrodes at any recording site, we increased the relative number of trials with long SOAs by using a flat distribution of SOAs in log2 space. This led to an approximately equal number of trials within each SOA octave, i.e., 250-500 ms, 500-1000 ms, 1000-2000 ms, etc. We have used the same log2 distribution in other EEG recordings and they did not seem to change the results in a meaningful way. (2) We adjusted the range of presented tonal frequencies to the preferred frequency at the recording site in question. However, because our focus was on understanding the microscopic correlate of the EEG signals which were recorded within a limited range of tonal frequencies (500 to 4000Hz) we mostly avoided frequencies that were too far outside of this range. For example, if the preferred frequency of the recording site was at 300 Hz, we would adjust the range of frequencies to include 300 Hz, but would position it at the lower edge to keep most of the range above 500 Hz. In the few cases where the preferred frequency was far outside the range of the EEG recordings, we ignored that preferred frequency and defaulted back to the EEG range.

### 4.7 Preprocessing of EEG data

Time-continuous raw EEG data was down-sampled from 5000 to 1000 Hz. The down-sampled data was then filtered using a 256 point non-causal digital low-pass FIR filter (*firws* function from EEG-lab toolbox, Blackman Window, high-frequency cutoff: 70 Hz, transition bandwidth: 21 Hz). The filtered data was then cut into short epochs around the onset of each sound (−150 to 750 ms at 500 Hz sampling rate and saved in a data format that was compatible with the statistics software R (R Development Core Team, 2009). A subtraction method was used to reduce ERP superposition for tones with short SOA (see Teichert et al 2016 for details).

### 4.8 Preprocessing of intracranial data

Multi-unit activity was estimated as the envelope of power in the frequency range between 500 and 3000 Hz. To that aim the data was first band-pass filtered in this frequency range, half-wave rectified, and then filtered with a 100 Hz low-pass. Because the duration of multi-unit responses was shorter, the subtraction method that was used for the EEG data was not necessary for the multi-unit responses. In some recording sessions, the raw intracranial data picked up two types of artifact: (1) electromagnetic interference caused by the speaker, (2) artifacts related to the presentation of the monkeylogic event triggers. Speaker artifacts could easily be regressed out since their exact shape and timing was known. The speaker artifact removal routine was only called if these speaker artifacts were actually present. Furthermore, the routine only removed potential artifacts if they exceeded a pre-defined threshold. Trigger artifacts could not be removed by regression because their exact shape was variable. Hence, we used an interpolation approach. Based on the magnitude of the artifact we either interpolated 6 or 20 time-points around the time of the trigger. The trigger artifact removal was applied to all data sets, regardless of whether or not there was clear visual evidence that they were affected. In most data sets, using or not using the trigger artifact removal routine had not discernible effect on the extracted multi-unit data. In addition to multi-unit data, we extracted single cell responses using the Blackrock Offline Spike Sorter (BOSS).

The resulting data were analyzed with a series of linear and non-linear models. A simple first-pass analysis was performed in a time-resolved manner. A secondary analysis was performed on responses averaged in specific time-windows. Lastly, we analyzed the data using a non-linear adaptation model.

### 4.9 Time-resolved linear model analysis

Averaged auditory evoked potentials were computed using in-house toolbox written in the statistics software R (R Development Core Team, 2009). EEG, LFP and MUA data were downsampled to 1000Hz and aligned to tone onset. To reduce the impact of movement artifacts, trials with peak-to-peak EEG amplitude above 450 μV were excluded. To reduce jaw muscle artifacts we rejected trials with high power in the 30-500 Hz range in susceptible EEG channels towards the edge of the implant. Data was aligned to tone onset and analyzed with a time-resolved linear model:

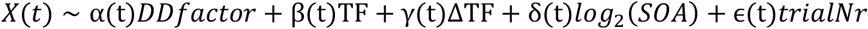

Here, X(t) corresponds to the time-resolved electrophysiological data being modeled. DDfactor is a categorical variable with four levels which arise as the interaction of the two key experimental manipulations: (1) the type of environment, random or regular, and (2) whether or not the current sound parameters (ISI and tonal frequency) were repeated or not. In the regular environment, a repeated sound is considered a standard, and a non-repeated sound a deviant. In the random environment, a non-repeated sound is considered a control. Repeated sounds in the random environment were not specifically analyzed in this context. (t) is a four-dimensional time-series, with each dimension corresponding to the activation profile of one of the four conditions described in DDfactor. TF is an 11-dimensional categorical variable that accounts for the differences in response magnitude as a function of the 11 different tonal frequencies. In case of the EEG data, it accounts for the inverted u-shape of EEG amplitude as a function of tonal frequency(Teichert, 2016). In case of the intracranial data, it accounts for the specific frequency tuning of the multi-unit in question. The variable ΔTF was introduced to account for a potential confound that was observed for the EEG data: changing tonal frequency by a certain amount from high to low typically leads to relatively larger responses than changing tonal frequency by the same amount from low to high. This effect could confound the effect that is being investigated here, namely the effect of the absolute value of the change in tonal frequency |ΔTF|. The fourth term is a first approximation for the effect of stimulus onset asynchrony on response magnitude. Finally, the fifth term was included to account for potential slow drift in response strength over the course of the entire experiment.

For the EEG data, this model was fit separately for each eeg-channel and each recording day. The parameter estimates (t) were then extracted and averaged across all recording sessions of a particular channel. Statistical differences between conditions were then assessed using time-resolved paired t-tests. For the intracranial data, the model was also fit separately for each recording contact and recording session. The parameter estimates (t) were then averaged across all recording contacts and recording sessions. Additional analyses separated the (t) either based on putative cortical layer or location on the superior temporal plane.

### 4.10 Scalar, non-time-resolved analysis

For the intracranial data, we followed up on the time-resolved analysis using a series of linear model analyses with activity averaged across pre-defined time-windows. These analyses used a very similar linear model. However, it excluded the main variable of interest DDfactor. Instead, the residuals of the model were extracted, z-scored and averaged according to DDfactor. This average z-scored data was then combined into a population analysis according to the latency of the pre-defined time-windows and the putative layer of the recording contact. These types of analysis allowed us to determine how SSA and DD depended on SOA and |ΔTF|.

An additional analysis was aimed at determining the percentage of single- and multi-units affected by several experimental factors such as tonal frequency, R, DD and SSA. This approach differed from the previous one along two lines. First, DDfactor was explicitly broken down into three distinct factors: response magnitude in the control condition served as the baseline. The factor DD was set to 1 on trials when a deviant was presented in the regular condition. The factor SSA was set to 1 when a tone was repeated, regardless of whether or not the repetition occurred in the regular or random condition. Finally, the factor REP was set to 1 on trials in the random condition when the tone was repeated. Together, these three factors explain the same variance as DDfactor, however, splitting them up allowed us to determine whether or not the parameter estimates were significantly different from zero, i.e., significantly different from the control condition. A unit with a significant positive main effect of DD will have stronger responses to the deviant relative to the control. A unit with a significant negative main effect of SSA will have weaker responses if a tone is repeated regardless of whether the repetition occurs in the random or regular condition. A unit with a significant effect of REP would indicate that the effect of SSA is different between the random and regular condition. We did not analyze or interpret the results of REP because they can either reflect differences caused by on average more repetitions in the regular condition, or an indication that unexpected repetition in the random condition are treated differently from expected repetitions in the regular condition.

### 4.11 Adaptation model

The STPSP model of SSA was designed to capture a several key aspects of short-term synaptic plasticity in the auditory system. The model assumes that responses of neurons in primary auditory cortex A1 are a linear function of excitatory post-synaptic potentials (**EPSP**) of thalamic inputs. The amplitude of the EPSPs is determined by thalamo-cortical synaptic weights, tonal frequency of the stimulus, and the receptive fields of the thalamic model neurons. In addition, each synapse is subject to short-term pre-synaptic depression such that synaptic strength is temporarily reduced in parallel with recent activity of the corresponding thalamic neuron. We refer to the ensuing model as the adaptation model. For each simulated tone, the adaptation model produces only one scalar value that can be interpreted, for example as total number of spikes in response to that tone, or the amplitude of a particular EEG component. This intermediate level of detail enables calculation of responses to hundreds of thousand of simulated tones within seconds, a prerequisite for fitting the model to the large data sets in this study using gradient descent approaches.

The adaptation model (Figure 8) consists of three main building blocks: (1) thalamic cells and their receptive fields; (2) short-term synaptic plasticity at thalamo-cortical synapses; (3) the receptive field architecture of pyramidal cells in primary auditory cortex. The receptive fields of thalamic neurons are modeled as a Guassians in log2 space. The parameter σ_Th_ describes the width of these receptive fields. Short-term synaptic plasticity at thalamo-cortical synapses is described by two parameters that quantify the fraction *U* of available vesicles *Q* that is expended for the processing of a single tone, and the recovery time constant *τ* with which vesicles are replenished back to baseline levels. The parameter *U* is closely related to release probability *u* in classic models of short-term synaptic plasticity. It is based on the assumption that a tone elicits a certain number of spikes *N*, and that each spike triggers the release of a fixed fraction of vesicles. The number of spikes *N* that a tone elicits in a thalamic cell is determined by the frequency of the tone, the preferred frequency of the thalamic cell and the width of its receptive field. If a tone elicits *N* spikes and each spike has a release probability of *u*=*15%*, the cumulative release probability, or ‘expended fraction’ *U* is given by:

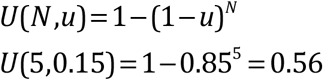

The parameter *U* describes the proportion of vesicles released across all *N* spikes elicited by a specific tone. Each thalamic cell was modeled to trigger N_max_ spikes to a tone of its preferred frequency. The number of spikes elicited by a tone of frequency *f* is given by:

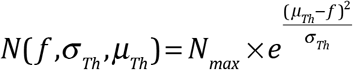

For the simulations *N_max_* was set to 5. The exact value of *N_max_* is only relevant if it is used to calculate the underlying value *u*, i.e., release probability per individual spike. Since this is not of interest here, any value *N_max_* would have provided the same result. The simulated EPSPs at each synapse are modeled to be proportional to the total number of vesicles released *R*, i.e, the product of *U* and *Q*:

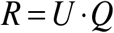

At the time of each tone presentation, the amount of available vesicles is reduced by the factor *U*:

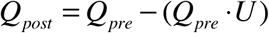

Between tones separated by an SOA of *y* seconds, the available vesicles *Q* recover exponentially back to baseline with the recovery time-constant *⊤*:

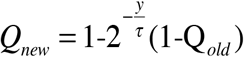

The structure of an A1 receptive field is described by the sets of synaptic weights to thalamic neurons. The adaptation model assumes two types of weights: 1) a ‘classical’ RF structure that samples densely from thalamic cells close to a certain preferred frequency, and a non-classic RF structure that samples uniformly from all thalamic cells regardless of their preferred frequency (pedestal). The classical RF structure is quantified by a Gaussian density function with width σ_A1_. The pedestal is defined by its amplitude relative to the peak synaptic strength of the classical RF component at the preferred frequency which was normalized to 1.

## 6 Author Contributions

TT conceived and designed the research. TT, KG and ZS performed the experiments. TT analyzed the data. TT wrote the manuscript. HJ built the intracranial recording setup. TT and HJ edited the manuscript.

